# Single-Cell ATAC-seq analysis via Network Refinement with peaks location information

**DOI:** 10.1101/2022.11.18.517159

**Authors:** Jiating Yu, Duanchen Sun, Zhichao Hou, Ling-Yun Wu

## Abstract

Single-cell ATAC-seq (scATAC-seq) data provided new insights into the elaboration of cellular heterogeneity and transcriptional regulation. However, scATAC-seq data posed challenges for data analysis because of its near binarization, high sparsity, and ultra-high dimensionality properties. Here we proposed a novel network diffusion-based method to comprehensively analyze scATAC-seq data, named **S**ingle-**C**ell **A**TAC-seq Analysis via Network **R**efinement with **P**eaks Location Information (SCARP). By modeling the prior probability of co-accessibility between adjacent peaks as a decreasing function of genomic distance, SCARP is the first scATAC-seq analysis method that utilizes the genomic information of peaks, which contributed to characterizing co-accessibility of peaks. SCARP used network to model the accessible relationships between cells and peaks, aggregated information with the diffusion method, and then performed dimensionality reduction to obtain low-dimensional cell embeddings as well as peak embeddings. We have demonstrated through sufficient experiments that SCARP facilitated superior analysis of scATAC-seq data. Specifically, SCARP exhibited outstanding cell clustering performance to better elucidate cell heterogeneity, and can be used to reveal new biologically significant cell subpopulations. SCARP was also instrumental in portraying co-accessibility relationships of accessible regions and providing new insight into transcriptional regulation, and those SCARP-derived genes were involved in some key KEGG pathways related to diseases. To sum up, our studies suggested that SCARP is a promising tool to comprehensively analyze the scATAC-seq data from a new perspective.

## Introduction

Chromatin accessibility is closely linked to the occurrence of transcriptional regulation, as accessible chromosome fragments often contain transcription factor (TF) binding sites and other important cis-regulatory elements, such as enhancers and promoters [1][2]. While single-cell transcriptome data can help us to construct gene regulatory networks from the level of transcripts and identify cellular diversity without bias, it is still challenging to reverse-engineer the mechanism of transcriptional regulation [2]. Fortunately, the development of single-cell epigenomic technology has brought new perspectives to solve this problem, such as single-cell assay for transposase-accessible chromatin using sequencing (scATAC-seq) [3] and single-cell combinatorial indexing assay for transposase accessible chromatin with sequencing (sci-ATAC-seq) [4][5], which can build chromatin accessibility profiles with single-cell resolution [6]. With the aid of scATAC-seq data, we were able to model the co-accessibility of chromosome segments from the genome level directly, which provides the basic conditions for transcriptional regulation to occur.

Compared with single-cell transcriptome data, scATAC-seq data exhibit more sparsity, higher dimension, and near-binarization properties, which pose many challenges to scATAC-based analysis. Because for a diploid genome, only one or two reads can be captured at each accessible chromatin site [6], resulting in the lack of sequencing data, i.e., the curse of ‘missingness’ in scATAC-seq [1][6]. In addition, accessible regions are usually called on the entire genome, so the features of scATAC-seq data can reach more than 100k dimensions or even higher. These properties make scRNA-seq-based algorithms less effective for analyzing scATAC-seq [7], although these algorithms are mathematically transferable since both genes and peaks can be understood as features to characterize cells.

Currently, various methods have been specially developed for scATAC-seq data to decipher their underlying information. For example, chromVAR [8] measures the accessibility gain or loss among peaks with the same motif or annotation and uses a bias-corrected deviation matrix for downstream analysis. However, the aggregation of similar peaks loses much information, making it less effective for cell clustering. SCALE [1] incorporates the variational autoencoder and Gaussian Mixture Model to extract latent features of cells. Nevertheless, it has numerous parameters and needs to be carefully tuned to get good performance [6]. Besides, cisTopic [2] uses the Latent Dirichlet Allocation model for co-optimal clustering of the cells and accessible regions, yet the collapsed Gibbs sampler makes it computationally slow. A recently proposed method, scAND [6], adopts the Katz index [9] diffusion method to alleviate the high sparsity in scATAC-seq data, and using the smoothed matrix as model input can achieve satisfactory results in their following analysis. However, scAND does not consider the differences between different chromosomes in the diffusion process and neglects the location associations of peaks.

In genetics studies such as recombination between two loci and linkage disequilibrium [10]–[13], the genetic map distance has been considered an essential factor. Genes on different chromosomes or distantly separated on the same chromosomes are assorted independently and described as physically unlinked [12], and the plausibility of modeling chromatin contacts as a decreasing function of genomic distances has been explained and verified [14]. Therefore, as scATAC-seq data give the accessibility of genome-wide segments in cells, incorporating genomic distance information of accessible fragments may be beneficial for the downstream data analyses, such as cell subpopulation identification and cis-regulatory network construction, which has not been considered in the state-of-the-art methods. In addition, it is also critical to exploit a superior network diffusion process that can appropriately aggregate the neighborhood information to deal with the sparseness and missingness problems in scATAC-seq data. Our previously developed Network Refinement (NR) [15] diffusion method, which is a degree-normalized version of the Katz index, has been shown to achieve better performances in a series of applications [15][16]. We, therefore, expect to see that NR also handles scATAC-seq data well, with the rationale being that: 1) The diffusion of accessibility relationships can compensate for missing information and depict similarities between nodes (i.e., cells and peaks) of the network; 2) Peaks accessible in too many cells should reduce their impact on diffusion, as their ability to portray cell similarity is overestimated (Methods).

In this study, we proposed a novel network-based method to comprehensively analyze scATAC-seq data, named **S**ingle-**C**ell **A**TAC-seq Analysis via Network **R**efinement with **P**eaks Location Information (SCARP). Specifically, SCARP takes genetic location information of peaks into account and globally diffuses the accessibility relationships using our previously developed Network Refinement diffusion method (Methods). The output matrix derived from SCARP can be further processed by the dimension reduction method to obtain low-dimensional embeddings of cells and peaks, which can benefit the downstream analyses such as the cells clustering and cis-regulatory relationships prediction (Fig. 1).

**Fig. 1.**
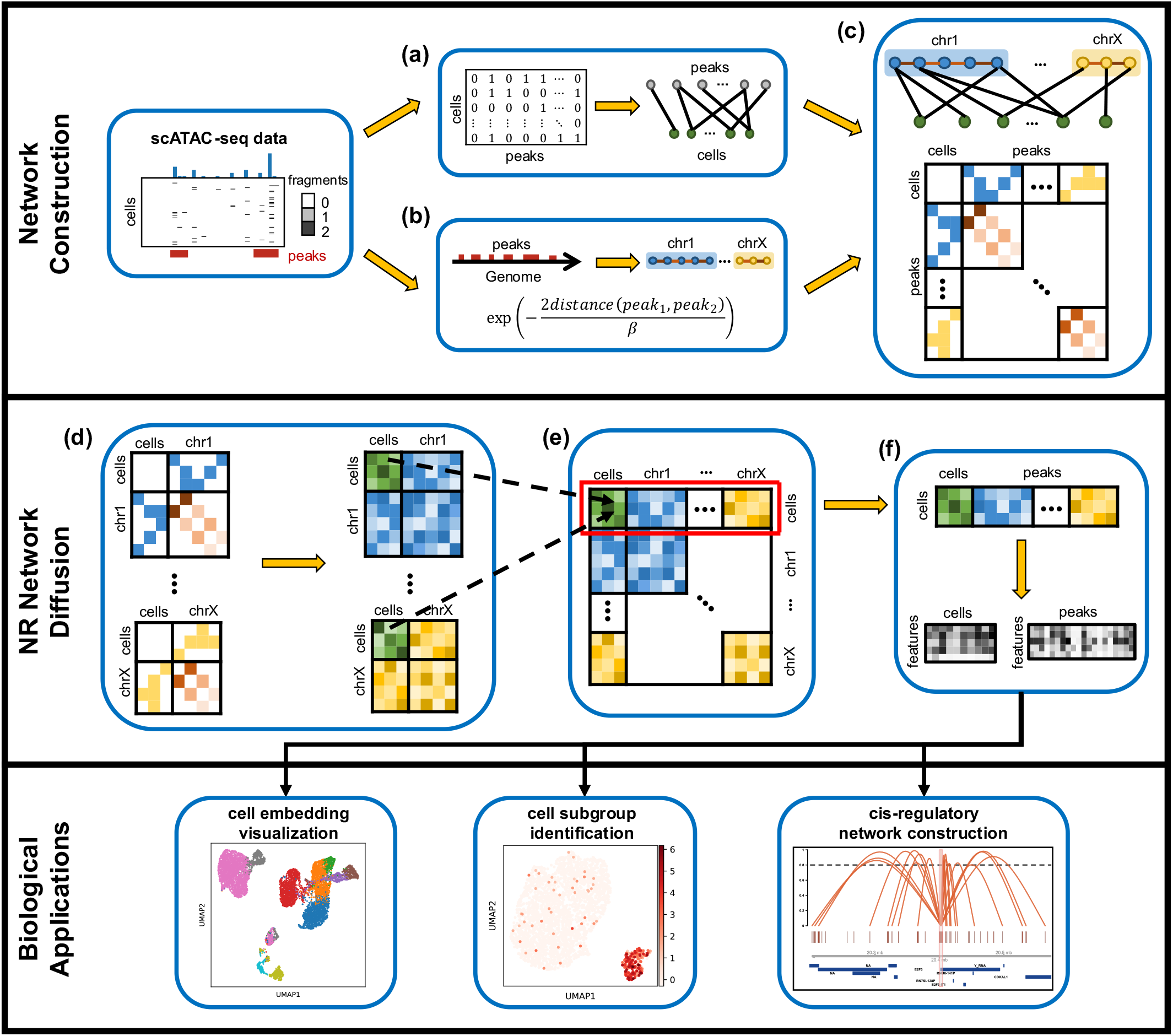
The workflow of SCARP. SCARP consists of two important modules, the first is constructing the network from scATAC-seq data, with the nodes being cells or peaks, the edges between cells and peaks reflecting accessibility, and the edges between peaks reflecting prior information about their genome position; the second is using NR diffusion to compensate for the effects of high sparsity of scATAC-seq data as well as characterize the similarity between cells. The output matrix of SCARP with the low-dimensional embedding of cells and peaks can further benefit the downstream analyses.

We have demonstrated through sufficient experiments that SCARP facilitates superior analysis of scATAC-seq data, including improving cell clustering performance to better elucidate cell heterogeneity and reveal new biologically significant cell subpopulations, and characterizing co-accessibility of genome segments to shed fresh light on transcriptional regulation. Our studies suggested that SCARP is a promising tool to comprehensively analyze the scATAC-seq data from a new perspective.

## Results

### Overview of SCARP workflow

We proposed a novel network diffusion-based method, SCARP, to comprehensively analyze the scATAC-seq data. The workflow of SCARP was shown in Fig. 1. First, SCARP constructed a bipartite network to model the accessible relationships between the cells and peaks (Fig. 1a). Second, prior edge weights of adjacent peaks based on their genome distance were employed to better capture the co-accessibility of adjacent peaks (Fig. 1b). Together they constituted the input network for the next step (Fig. 1c). Third, the NR diffusion processes [15] were performed on the subnetworks corresponding to different chromosomes separately to obtain dense matrixes that can reflect accessibility similarities between the cells and peaks, as well as possible cell-peak accessible relationships (Fig. 1d). After this, diffusion matrixes obtained from different chromosomes were spliced together and the cell-cell similarities were integrated (Fig. 1e). Finally, dimensionality reduction technique was performed to map cells and peaks into low-dimensional space, and these representations were used for downstream analyses, such as cells clustering and construction of cis-regulatory networks (Fig. 1f).

### SCARP exhibited superior cell clustering performance on benchmarking scATAC-seq datasets

We selected nine benchmarking scATAC-seq datasets with reference cell type annotations. For these datasets, the number of peaks ranges from 7,000 to 140,000 with different sparsity levels (Fig. 2a and Supplementary Table 1). We compared SCARP with six state-of-the-art methods (original, DCA [17], MAGIC [18], PCA, cisTopic [2], and scAND [6]) as well as SCARP without using prior weights of adjacency peaks (denoted as SCARP*). Notably, DCA and MAGIC were especially designed for scRNA-seq analysis (Supplementary Materials). The running time of SCARP is advantageous when compared with these methods (Fig. 2b and Supplementary Table 3). Using blood2K dataset as an example, we found stronger consistency between the identified cell clusters using SCARP cell embeddings and the annotated cell types when compared with other candidate methods (Fig. 2c and Supplementary Figs. 1-2). The SCARP-derived low dimensional visualization of cells also exhibits more explicit cell type boundaries and a consistent developmental trajectory with known FACS-sorted population [6][19] (Fig. 2d and Supplementary Figs. 3-6). For all benchmarking datasets, SCARP achieved the top-ranked clustering performances with different evaluation metrics (Fig. 2e and Supplementary Fig. 7). It is noteworthy that SCARP* ranked second, meaning that both NR diffusion and prior weights play essential roles in obtaining better low-dimensional cell representations. Together with the observations that SCARP had the best averaged clustering performances regarding different metrics (Fig. 2f, g), all these pieces of evidence comprehensively demonstrated the superiority of SCARP over other methods in cell feature extraction.

**Fig. 2.**
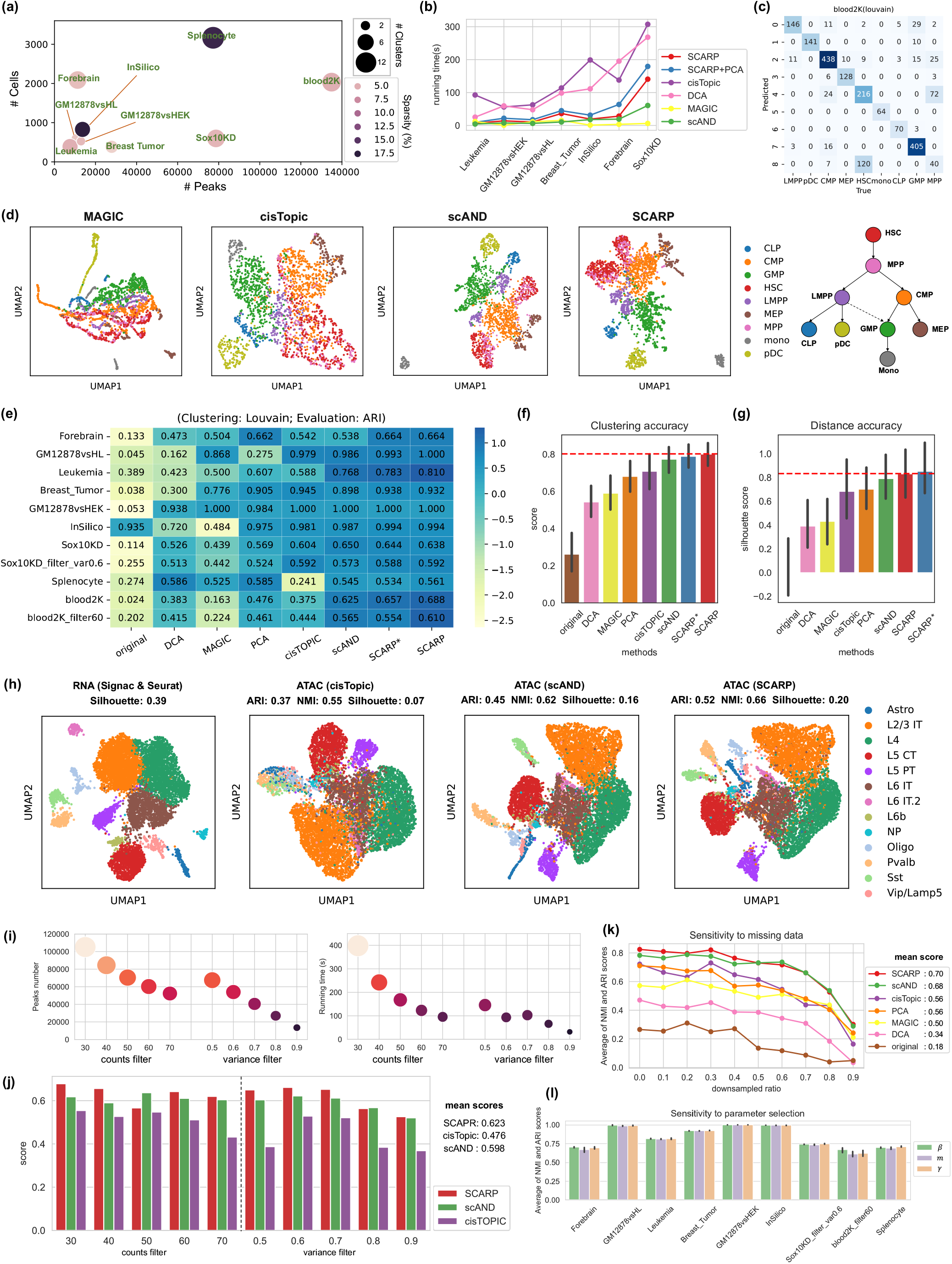
Cell clustering performance on benchmarking scATAC-seq datasets. **(a)** Data presentation. Each dot represents a scATAC-seq dataset, with the x-axis (y-axis) showing its number of peaks (cells), the size of the dots showing the number of annotated cell types, and the color of the dots showing the data sparsity. **(b)** Running times (y-axis) of various methods on seven benchmarking datasets (x-axis). One method is shown by one line (color). **(c)** The confusion matrix of SCARP-derived cell embeddings clustered by Louvain (rows) and the annotated cell types (columns) on the blood2K dataset. The matrix elements represent the number of overlapping cells in a predicted cluster and a real cluster. **(d)** The UMAP plots of four methods MAGIC, cisTopic, scAND, and SCARP on the blood2K dataset with 2034 cells, colored by the annotated cell types. The rightest legend was the known FACS-sorted population of origin, which shows the process of cell development and differentiation. **(e)** The cell clustering results of eight methods (x-axis) on eleven datasets (y-axis) under ARI evaluation. Colors represent the Z-scores calculated from ARI score of the corresponding dataset. **(f)** Cell clustering accuracy (y-axis) of various methods (x-axis). Scores were calculated by averaging the ARI scores and NMI scores for a method over all datasets. **(g)** Cell distance accuracy (y-axis) of various methods (x-axis). Scores were calculated by averaging the silhouette coefficient scores for a method over all datasets. **(h)** Four UMAP visualizations showing the cell clustering results of Seurat & Signac based on scRNA-seq data, as well as cisTopic, scAND, and SCARP based on scATAC-seq data, and their clustering accuracy scores (title). The plots were colored by the annotated cell types obtained by Seurat. **(i)**The x-axis represents different peak filtering strategies, and the y-axis on the left panel represents number of peaks in the blood2K dataset, the y-axis on the right panel represents running time of SCARP, **(j)** Barplots of cell clustering performance (y-axis) for SCARP, scAND and cisTopic under different strategies of peaks filtering (x-axis). Scores (y-axis) were calculated by averaging the ARI score and NMI score for one method. Mean score (right legend) for each method was calculated by averaging the scores for all peak filtering strategies. **(k)** The average of NMI and ARI scores (y-axis) of various methods on the Leukemia dataset with different downsampling ratios (x-axis). **(l)** The average of NMI and ARI scores (y-axis) when taking different values of parameters *β, m*, and *γ* on various scATAC-seq dataset (x-axis). Error bars represent the standard deviation (SD) of the scores.

Since cell type annotations based on RNA-seq data are more readily available, we further investigate whether the scATAC-derived cell clustering is consistent with the scRNA-based results, using the SNARE-seq data, which is paired multi-omics data that simultaneously profile gene expressions and chromatin accessibilities [20]. We used the cell clustering results on RNA-seq data processed by Seurat [21] and Signac [22] as a reference, and compared their consistency with those of SCARP, scAND, and cisTopic on scATAC-seq data. We observed that SCARP yielded higher cell clustering concordance between paired multi-omics data (Fig. 2h).

We next explored the robustness of SCARP in dealing with different peak numbers, missing data, and different model parameters. Since the blood2K dataset has the maximum number of peaks, we used it to test the impact of peak filtering to the performance of SCARP. We found that the number of peaks and the running time of SCARP decreased as the filtering level increased under both count-based and variance-based peaks filtering strategies (Fig. 2i). In contrast, the accuracy of cell clustering of SCARP had not been much affected, and the cell clustering performances were still better than that of two other competitive methods for most filtering levels (Fig. 2j). Besides, we randomly down-sampled the counts in Leukemia dataset, which has the minimum number of peaks, to simulate the missing values in scATAC-seq data. We observed that our method was not sensitive to missing data for different down-sampling ratios and got the best performance overall (Fig. 2k and Supplementary Fig. 8). Furthermore, SCARP has three model parameters (*β*, *m*, and *γ;* Methods). We found that SCARP is robust to the parameter selection since the cell clustering results only slightly fluctuated (Fig. 2l and Supplementary Figs. 9-11).

Taking together, SCARP is a fast and robust method to extract informative low-dimensional cell embeddings and has an outstanding performance of cell clustering compared with all the other methods.

### SCARP identified a new CD14 monocyte subpopulation characterized with high differentiation activities

We applied SCARP to a 10X multiome peripheral blood mononuclear cell (PBMC) dataset [23], where scRNA-seq and scATAC-seq data can be acquired simultaneously. The cell clusters generated using SCARP-derived cell embeddings were well-separated between different cell types (Fig. 3a). Interestingly, we observed that the cells originally annotated as CD14 monocytes were divided into two groups (denoted as cluster 1 and cluster 2) in UMAP visualization (Fig. 3a and Supplementary Fig. 12), indicating the existence of possible new subpopulations of CD14 monocytes. Differentially expression analysis showed that cluster 2 monocytes were characterized by the overexpression of *BCL11B, CD247*, and some interleukin-related genes (Fig. 3b and Fig. 3d). Several marker peaks were also identified corresponding to each monocyte subpopulations (Fig. 3c, Fig. 3e and Supplementary Figs. 13-14).

**Fig. 3.**
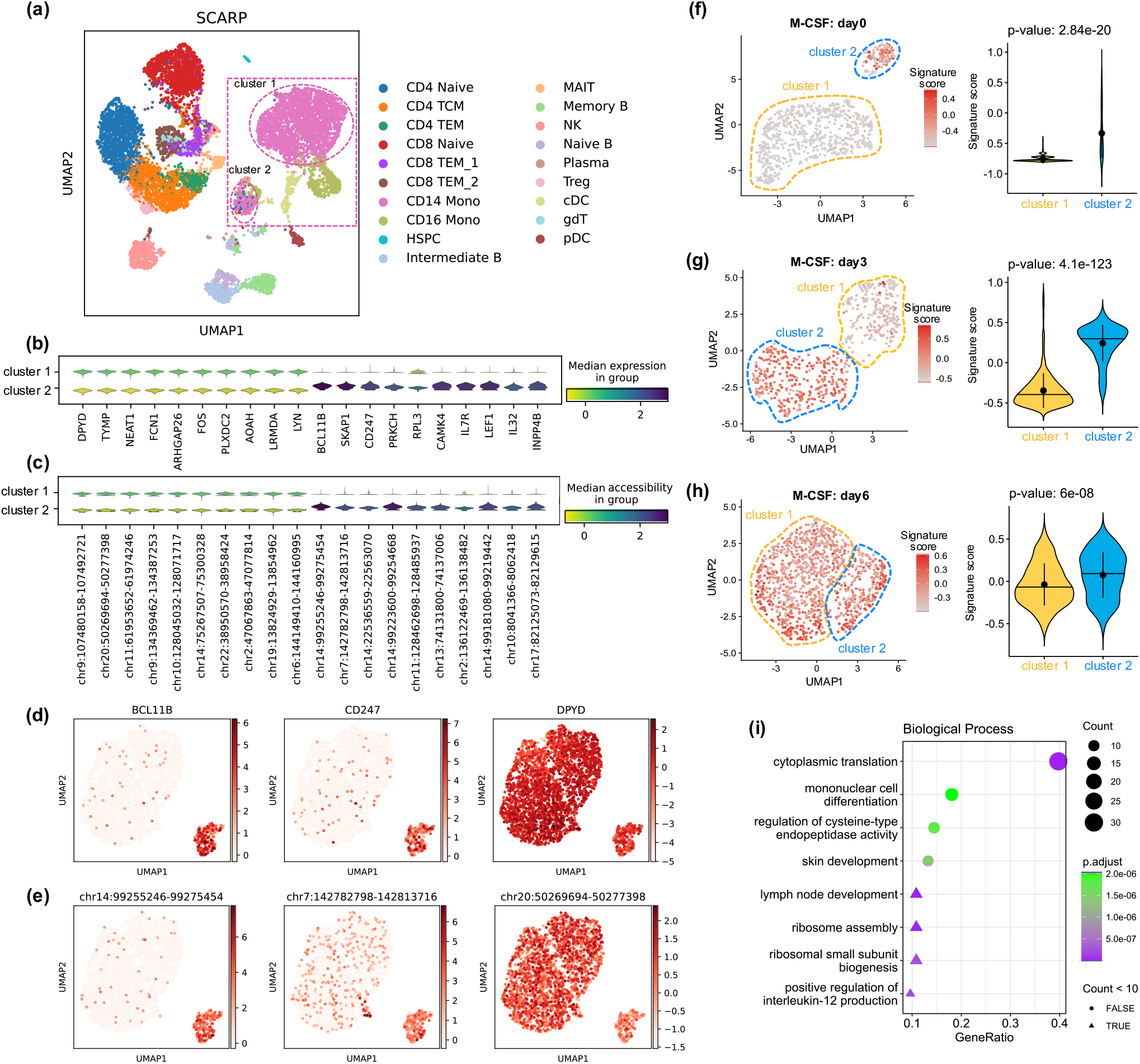
SCARP identified a new CD14 monocyte subpopulation. **(a)** UMAP visualization of SCARP-derived cell embeddings colored by the cell types annotated by Seurat. Those CD14 monocytes were framed in pink, and two cell subgroups were detected, denoted as cluster 1 and cluster 2. **(b)** Violin plots showing the expression levels of the top 10 marker genes (x-axis) for two cell subpopulations (y-axis). **(c)** Violin plots showing the accessibility levels of the top 10 marker peaks (x-axis) for two cell subpopulations (y-axis). **(d)** The UMAP visualizations of cells colored by the expressions of cluster 2 marker genes *BCL11B, CD247*, and the expressions of cluster 1 marker gene *DPYD*. **(e)** The UMAP visualizations of cells colored by the accessibility of cluster 2 marker peaks chr14:99255246-99275454 and chr7:142782798-142813716, and the accessibility of cluster 1 marker peak chr20:50269694-50277398. (f)(g)(h): The UMAP visualizations of cells in external M-CSF scRNA-seq dataset, colored by gene signature scores for CD14 monocytes (donor 1) under M-CSF stimulation at day 0 **(f)**, day 3 **(g)** and day 6 **(h)**, along with the violin plots showing the difference between the two cell clusters. **(i)** Top eight biological processes that gene signatures are involved in, using all genes in this scRNA-seq dataset as the background gene set. The y-axis represents the name of KEGG pathway, and the x-axis represents the ratio of genes enriched in a pathway. The number of genes enriched in a pathway is related to the size of the circle (gene count less than 10) or triangle (gene count greater than 10), and the color represents the corresponding adjusted p-value.

To ascertain whether the newly identified CD14 monocyte subpopulation is biologically meaningful, we built a CD14 monocyte signature using the top 50 marker genes of cluster 2 monocytes. After this, we applied our CD14 monocyte signature to a time-series scRNA-seq dataset of human CD14 monocytes, where CD14 monocytes were stimulated by macrophage colony-stimulating factor (M-CSF) at days 0, 3, and 6 [24]. We observed that only 15% of monocytes were clustered together, and 25% of them had relatively higher CD14 monocyte signature scores at day 0 (Fig. 3f, Student’s *t*-test *P* = 2.84e-20). This finding became more apparent at day 3, with approximately half of the monocytes had significantly higher signature scores (Fig. 3g, Student’s *t*-test *P* = 4.1e-123). With the continuous stimulation of M-CSF, almost all cells (86%) exhibited high signature scores, and the difference between the two clusters was less significant (Fig. 3h, Student’s *t*-test *P* = 6e-08). These observations can be reproduced using other replicates (Supplementary Figs. 15-16). Besides, we also identified the new CD14 monocyte subpopulation in another CD14 monocyte scRNA-seq dataset with higher signature scores (Supplementary Fig.17).

Since monocytes are precursors to macrophages and can differentiate into dendritic cells, we hypothesized that the signature scores can serve as differentiation activities and the monocytes with high signature scores are prone to differentiate into other cells. We thereby conducted GO functional enrichment analysis using the signature genes. We found that these genes were associated with mononuclear cell differentiation and its related processes, such as T cell differentiation [25] and immune response-activating signal transduction [26] (Fig. 3i and Supplementary Fig. 17). These results confirmed the ability of our gene signature to identify CD14 monocyte subtypes.

In conclusion, we demonstrated that the SCARP-derived low-dimensional representation of cells can refine cell types and reveal biologically meaningful cell subtypes.

### SCARP discovered key factors of melanoma progression by portraying co-accessibility relationships of peaks

Except for the low-dimensional cell embeddings, SCARP can also generate biologically meaningful peak embeddings. In this experiment, we want to investigate whether the SCARP-derived peak embeddings can uncover some potential regulatory relationships in melanoma disease progression. To achieve this, we applied SCARP on a *SOX10* knockdown time series scATAC-seq dataset (SOX10KD) [2], which records changes in the accessibility of peaks at 0, 24, 48, and 72 hours after *SOX10* knockdown in two patients (MM057 and MM087).

We first annotated two regions related to *SOX10*, i.e., the promoter (chr22:38380499-38380726) and 3’UTR (chr22:38364975-38365257) of it (Supplementary Fig. 18). Since the promoter initiates gene transcription while 3’UTR can decrease the expression of the corresponding gene [27] [28], it is reasonable to see that the accessibility of *SOX10* promoter increases and 3’UTR decreases as knockdown time passes. By calculating the correlations between peaks’ low-dimensional representations, we obtained the top 10 co-accessible regions with the highest correlation to each of the *SOX10* promoter and 3’UTR. These loci have the more similar time-varying trend as the *SOX10* promoter (3’UTR) than that found by using original data, indicating that SCARP can well characterize the co-accessibility of accessible segments (Fig. 4a and Supplementary Fig. 25a). To better interpret these co-accessible regions, we obtained the corresponding genes using the promoter regions (Supplementary Fig. 19; Methods) and selected the top 50 genes with the most co-accessible relationships with *SOX10*. By constructing a gene regulatory network (GRN) between these genes, we found abundant database-supported regulatory relationships (Fig. 4c). Notably, SCARP owned the largest number of curated edges and the second number of verified genes in the constructed GRNs compared to other candidate methods, such as cisTopic and scAND (Fig. 4b).

**Fig. 4.**
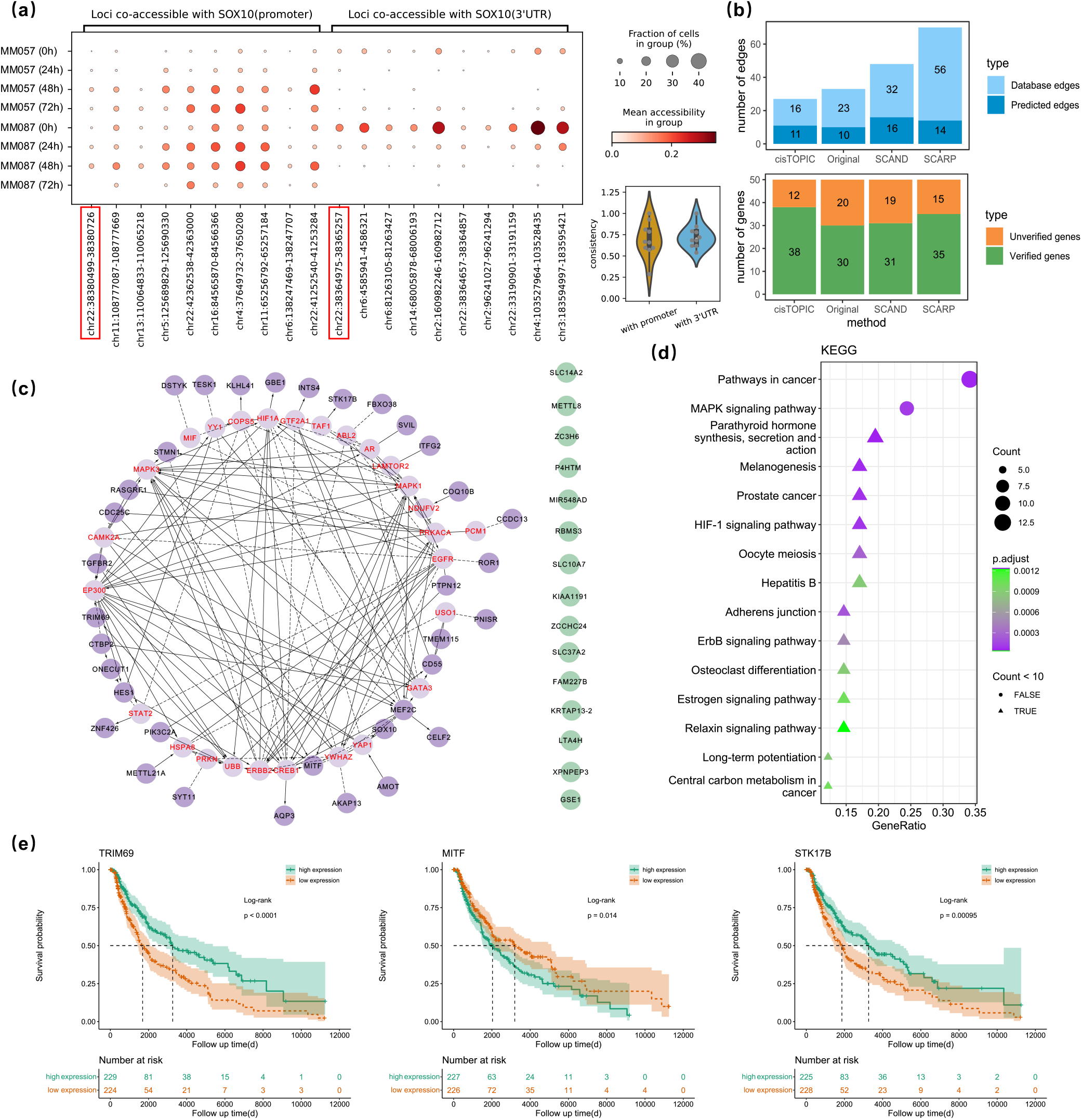
SCARP portrayed the co-accessibility of peaks on SOX10KD scATAC-seq dataset. **(a)** Changes in accessibility for those loci (x-axis) that were highly correlated with *SOX10* promoter and 3’UTR as knockdown time walked by (y-axis). The *SOX10* promoter and 3’UTR were framed in red. The size of the dots showing the fraction of cells in cell groups, and the color of the dots showing the mean accessibility in cell groups. The violin plot at the right bottom showed the consistency of the trend for the top ten peaks coaccessible with the promoter (3’UTR) of SOX10, with consistency calculated from the correlation of mean accessibility within eight cell groups. **(b)** For each of the differential gene sets (50 genes) obtained by four methods (x-axis), we compared the number of genes (y-axis of lower panel) that were verified (green bars) as directly or indirectly related by the database, and the number of genes that were not verified (orange bars), as well as the number of edges (y-axis of upper panel) in the corresponding GRNs, including predicted edges (dark blue), i.e., those with a functional interaction (FI) type of “predicted”, and database edges (light blue), i.e., those with a FI type other than “predicted”, such as “activate” and “inhibited by”. **(c)** The transcriptional regulatory relationships validated by the Reactome database. Solid lines indicate experimentally validated regulatory relationships, while dashed lines indicate database-predicted regulatory relationships. The dark purple nodes, light purple nodes, and green nodes represent the SCARP-derived genes, linker genes, and unvalidated genes, respectively. **(d)** Results of KEGG pathway analysis. The legends are the same as in Fig. 3i. **(e)** The results of survival analysis on genes *TRIM69, MITF*, and *STK17B*.

Among the genes in SCARP-derived GRN, numerous genes are strongly associated with melanoma disease. For example, the upregulations of *STMN1* and *ROR1* could contribute to tumor migration and proliferation [29][30][31]. *LTA4H* regulates the cell cycle process and can lead to skin carcinogenesis by its overexpression [32]. Notably, we uniquely identified that *HIF1A*, which is known for promoting cancer progression and causing therapy resistance [33][34], interacts with the key master regulator of melanocyte development and melanoma deterioration *MITF* [35][36] (Fig. 4c). Numerous studies have reported that hypoxia is closely related to the invasiveness, angiogenesis, and therapy response in melanoma [33][37][38]. In studies of its pathogenic mechanism, *MITF* has been shown to stimulate the transcriptional activity of *HIF1A* to supply oxygen to cancer cells [39][40].

We further conducted the functional enrichment analysis using the genes in SCARP-derived GRN. As a result, we found many melanoma progression-related KEGG pathways, e.g., melanogenesis [41][42], MAPK signaling pathways [43][44], and Adherens junction [45][46] (Fig. 4d and Supplementary Fig. 21). Notably, the HIF-1 signaling pathway was again exclusively identified by SCARP (Fig. 4d and Supplementary Figs. 23-25) [47][48]. In addition, GO enrichment categories contained several melanoma-related terms, such as muscle tissue development [49] and membrane raft [50] (Supplementary Fig. 22). Furthermore, we performed survival analysis using the genes in SCARP-derived GRN and found that many genes’ abnormal expressions were associated with the poor prognoses, such as *TRIM69, MITF, STK17B* (Fig. 4e and Supplementary Fig. 26). In particular, the low expression of gene *STK17B*, which has been shown in the literature to play an important role in immune infiltration, was associated with skin cancer, making it a key biomarker for the diagnosis of melanoma disease [51].

### SCARP predicted reliable cis-regulatory interactions supported by external evidence

We next explored the potential of the SCARP-derived peak embeddings and investigated their cis-regulatory relationships. We first defined the co-accessibility score of two peaks as the cosine similarity of their low-dimensional peak embeddings. This co-accessibility score can serve as the confidence of the corresponding cis-regulatory interactions. To assess the reliability of this score, we use external evidence to validate them, such as promoter-capture Hi-C (PCHi-C) [52][53] and ChIP-seq data [54]. Specifically, we selected those peaks belonging to promoter regions of certain genes and focused on the regulatory relationships between these peaks (i.e., promoters) with other peaks.

As PCHi-C data depicted whether there were physical interactions between the promoters (baits) and the entire genome (other ends), we can treat whether SCARP-derived cis-regulatory relationships were validated by PCHi-C data as a binary classification problem and measure its accuracy by calculating AUROC score (Methods). The result shows a significant difference between PCHi-C-validated and unvalidated SCARP-derived cis-regulatory scores (Fig. 5a, Student’s *t*-test *P*=1.31e-230), and the receiver operating characteristic (ROC) curve demonstrates the superiority of SCARP over other methods (Fig. 5b).

**Fig. 5.**
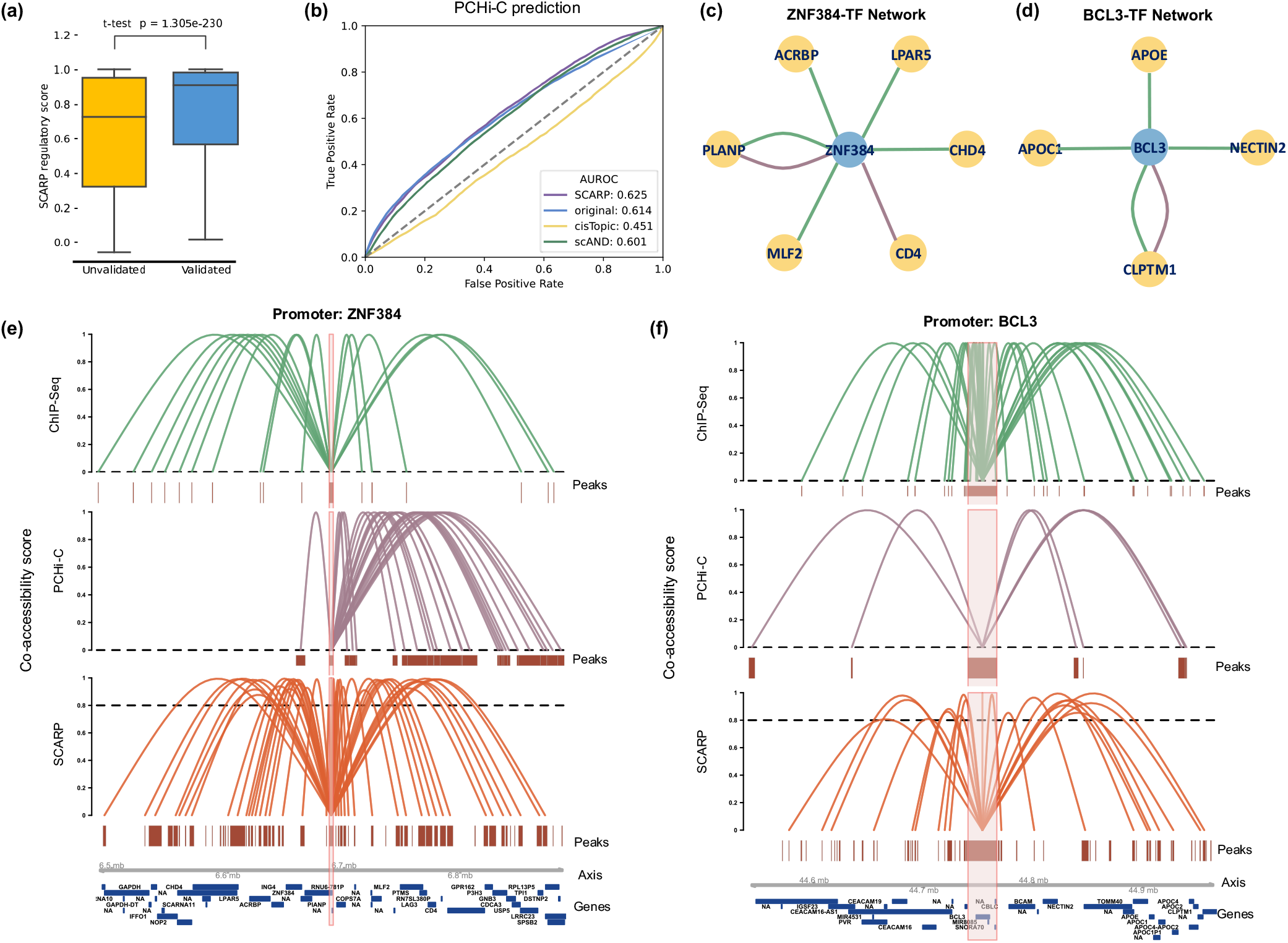
SCARP-derived cis-regulatory interactions are significant and validated by external evidence. **(a)** The boxplot showing the difference between PCHi-C validated (blue) and unvalidated (yellow) SCARP-derived transcriptional regulatory scores (y-axis), followed by t-test to test the difference. **(b)** ROC curve of various methods, treating whether SCARP-derived transcriptional regulatory relationships were validated by PCHi-C data as a binary classification problem. (c), (d), SCARP-derived TF-regulon networks of genes *ZNF384* **(c)** and *BCL3* **(d).** The green and purple edges mean those interactions are also supported by ChIP-seq or PCHi-C data, respectively. The blue and yellow nodes mean the TF genes and target genes respectively. (e), (f), Visualizations of co-accessibility scores (y-axis) of SCARP-derived cis-regulatory interactions (orange), PCHi-C evidence (purple), and CHIP-seq evidence (green) around the genes **(e)** *ZNF384* locus (x-axis) and **(f)** *BCL3* locus (x-axis).

Besides, we also used ChIP-seq data of human TF to validate SCARP-derived cis-regulatory relationships [54], which gives the experimentally verified interaction between proteins and DNA to help identify TF binding sites. To use evidence from both PCHi-C and ChIP-seq data, we selected those peaks that were both promoter regions and corresponded to a certain TF to see if the interaction with this region could be verified (Methods; Supplementary Fig. 27). As a demonstration example, we plotted the cis-regulatory interactions in the promoter region of gene *ZNF384* (Fig. 5c and Fig. 5e), which is a human TF associated with B-cell acute lymphoblastic leukemia (B-ALL) disease [55][56]. Visually we see that many of the SCARP-derived interactions were validated by either Chip-seq or PCHi-C data. We further extracted the *ZNF384* regulon network by mapping the SCARP-derived peaks to gene level, as shown in Fig. 5c, and those genes (yellow nodes) interacting with *ZNF384* can be regarded as its target genes, since they were proven to have its binding sites (green edges). Some literature evidence can also be found to verify this result: *MLF2* is a key factor associated with leukemia disease [57], and *CHD4* is stated as a potential therapeutic target of acute myeloid leukemia [58][59]. Another example of the promoter region of gene *BCL3* shown in Fig. 5d and Fig. 5f exhibited similar results. *BCL3* is also related to chronic lymphocytic leukemia and B-cell malignancies [60][61], and the genes interacting with it were supported by external data and literature [62][63][64]. More examples can be found in Supplementary Fig. 27.

## Discussion

Single-cell ATAC-seq data established genome-wide chromatin accessibility profiles with single-cell resolution that could complement single-cell transcriptome data, and together they provided new insights into transcriptional regulatory mechanisms. The accessible regions of scATAC-seq data, which can be seen as cell features, exhibited many different properties from genes of scRNA-seq data, such as higher sparsity, higher dimension, and near-binarization [1][6]. Thus, even both peaks and genes can be used to characterize cellular heterogeneity, and many scRNA-seq-based algorithms are mathematically transferable to analyze scATAC-seq, we should not be limited to previous attempts and ideas for dealing with scRNA-seq data given the different nature of the two features; instead, we should develop new approaches that focus on the specific properties and information of the scATAC-seq data itself and elaborate on cellular heterogeneity from new perspectives.

Although the severe signal deficiencies and high sparsity of scATAC-seq data are well known, the importance of filling in missing data and denoising was not given enough attention in some commonly used scATAC methods [65]. In this study, we proposed a network diffusion-based computational approach to comprehensively analyze scATAC-seq data. Considering the high sparsity of the data, it is very appropriate and natural to use the diffusion method for aggregating the neighborhood information between two nodes. Our previously developed NR diffusion method [15] has achieved outstanding success in many other applications, such as network denoising [15] and link prediction [16], and in this study it also proved to have excellent performance in handling scATAC-seq data.

In addition, to mine and exploit the unique information of scATAC-seq data, we have borrowed ideas from many genomic studies and used the genetic map distance as an important priori factor in the analysis of epigenomic data. To further test the soundness of this hypothesis, we measured the co-accessibility likelihood between two peaks using the p-value of Fisher’s exact test, and then investigated its relevance to genomic distance of peaks. The results shown in Supplementary Fig. 28a confirm that peaks closer in the genome are more likely to be co-accessible. The peaks with network weights higher (or lower) than expected after diffusion show a certain pattern on the genome (Supplementary Fig. 28b-e), i.e., the posterior probabilities of co-accessibility in some regions are higher (or lower) than that predicted solely based on genomic distance. The enrichment of the different region types within the SCARP-derived low-dimensional peak embedding features also supported that there were clear differences in accessibility between the different region types, which were closely related to their regulatory functions (Supplementary Fig. 20). Furthermore, the results of our experiments supported that the use of prior edge weights between adjacent peaks facilitate SCARP to mine more valuable information from scATAC-seq data, and also helped to obtain better downstream analysis performance.

We have demonstrated through sufficient experiments that SCARP comprehensively promotes superior analysis of scATAC-seq data compared to other state-of-the-art methods, such as scAND [6] and cisTopic [2]. Of all methods evaluated, SCARP had the best averaged clustering performances on benchmarking scATAC-seq data, and was robust to different peaks filter strategies and parameter selections. The running time of SCARP is also advantageous since we used some appropriate tricks to reduce the time cost. For example, to make those peaks on the same chromosome better learn from each other, we performed NR diffusion on subgraphs in parallel and spliced them back together.

Apart from the benchmarking dataset, we have designed three innovative case studies to explore what kind of analyses could be done with the scATAC-seq data to contribute to our further understanding of biological regulatory mechanisms. Specifically, we used SCARP-derived cell embeddings to identify a new CD14 monocyte subpopulation. Differential analysis between the two cell clusters obtained by SCARP identified a CD14 monocyte signature, which was later proved to be associated with mononuclear cell differentiation. We also applied SCARP on a melanoma-related scATAC-seq data to uncover some potential regulatory relationships in melanoma disease progression. Our analyses discovered several key factors of melanoma using SCARP-derived peak embeddings, such as genes *MITF, HIF1A*, and *STMN1*. Altogether, this showed how SCARP can be used to reveal disease mechanism by portraying co-accessibility relationships of peaks. Furthermore, the comparison of co-accessibility scores between SCARP-derived cis-regulatory interactions of peaks with external evidence, such as PCHi-C and CHIP-seq data, demonstrated the reliability of SCARP’s peak low-dimensional embeddings.

In the future, we expect to explore more unique properties of the scATAC-seq data to better handle it, and consider doing joint analyses with other multi-omics data, such as data alignment and integration.

## Methods

### SCARP workflow

Given scATAC-seq data matrix *A*_*c*×*p*_ with *c* cells and *p* peaks, we first binarized *A* to better cater to its biological interpretation of depicting accessibility. Specifically, we set all values greater than 1 to 1, making *A* a Boolean matrix with entries from the Boolean domain {0, 1}, reflecting whether a cell is accessible on a certain peak. Then a symmetric square root graph normalization 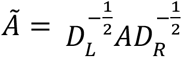 was applied to eliminate the library size differences between cells and peaks, where *D_L_* and *D_R_* are diagonal degree matrixes with (*D_L_*)_*ii*_ = ∑*_j_A_ij_* and (*D_R_*)_*jj*_ = ∑_*i*_ *A_ij_* [6].

After the pre-processing step, SCARP constructed a bipartite network with an adjacency matrix *B*_*N*×*N*_ to model the accessible relationship between the cells and peaks, where *N* = *c* + *p* is the number of nodes of the network, i.e., the peaks and cells are all treated as nodes of the network. To incorporate genetic location information for a better depiction of the co-accessibility of adjacent peaks, we computed the prior weights between adjacent peaks using the negative Haldane’s genetic map function. That is to say, we can now represent the network as:

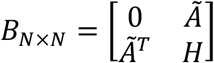

where *H* is a sub-diagonal matrix of size *p* × *p*, whose element represents the prior weight between the adjacent peaks in the same chromosome.

NR diffusion method was then applied to compute accessibility similarities between the cells and peaks, as well as predict possible cell-peak accessible relationships. Considering that those peaks on the same chromosome can better learn from each other, the NR diffusion process was calculated on the subnetwork of *B*, which is induced from nodes of cells and those peaks on the same chromosome, and then spliced back together. Specifically, the graph *B* was divided into a number of subgraphs 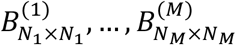, where *N_k_* = *c* + *p_k_* and *p_k_* is the number of peaks in the subgraph *B*^(*k*)^, and NR was performed on *B*^(*k*)^ to get a diffused matrix. This diffusion process can be performed in parallel, which reduces the time cost. Notably, when splitting peaks, we divided them according to whether they were on the same chromosome or not by default. However, given the sparsity of the scATAC-seq data, when the number of peaks is too small, we merged the adjacent chromosomes, and set the threshold parameter *γ* (default as 3000) to determine whether we need to merge the adjacent chromosomes.

Notice that after parallel diffusion, we got *M* dense matrixes, 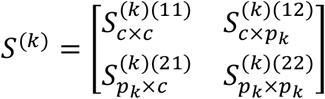, each of which has a submatrix of size *c* × *c*, 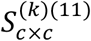, representing the cell similarity calculated based on the peaks of the current subgraph, and when splicing them back into matrix *D*_*N*×*N*_:

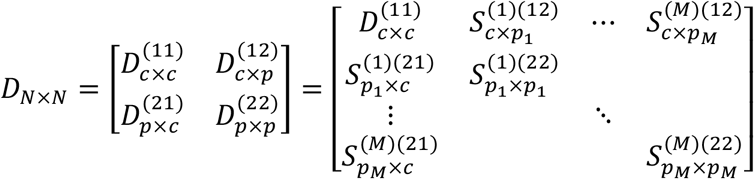

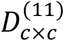 was the average over *M* matrixes 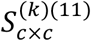 (Fig. 1).

Finally, to get cell and peak embeddings, the dimensional reduction method, such as UCPCA, was performed on the matrix 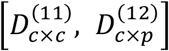 and 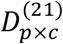 separately. The number of kept components was determined based on the proportion of variance that can be explained, as detailed below.

### Prior weights between adjacent peaks

The prior edge weights between the adjacent peaks in this study were calculated using the negative Haldane’s genetic map function, motivated by the traditional use of gene map function [10] to estimate the relation connecting recombination fractions *r* and genomic distances *d*:

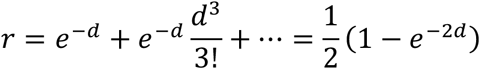

We assumed that the probability of chromosome segments being co-accessible should be a decreasing function of its genomic distance, as opposed to the probability of recombination [52][54]. Specifically, the distance between two peaks peak_1_ = [*s*_1_, *e*_1_] and peak_2_ = [*s*_2_, *e*_2_] (any two peaks do not intersect with each other) was *d*(peak_1_, peak_2_) = max (*s*_2_ – *e*_1_, *s*_1_, – *e*_2_), where *s_i_* stands for the start position and *e_i_* stands for the end position on the genome, and the prior weight of the two peaks was computed as:

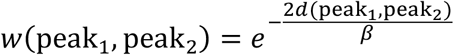

here we set a parameter *β* (default as 5000) to control the extent to which prior edge weight decays as peak distance increases.

### NR diffusion method

We used our previously proposed method, Network Refinement (NR) [15], to cope with the sparsity of scATAC-seq data. Specifically, NR takes the adjacency matrix of a network as input, uses a diffusion process defined by a random walk on the graph to enhance the self-organization properties of complex networks, and outputs a network with adjusted edge weights. The matrix obtained by NR reflects both the similarity of cells and peaks, and gives a confidence score for the possible cell-peak accessible relationship.

The graph operator *F_m_* of NR is defined based on three operators: *f_m_*, *g*, and *h*. The operator *f_m_* transforms a transition matrix *P* to another by adding the probability of all paths of different lengths joining two nodes, with a smaller weight coefficient 1/*m^k^* for a longer path of length *k*:

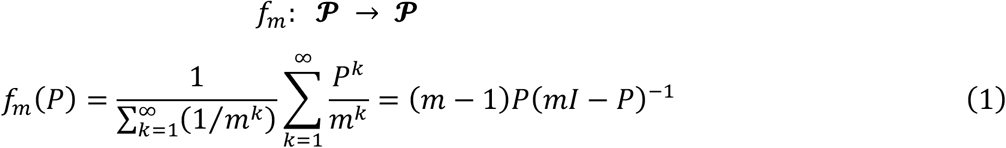

where ∑_*k*_(1/*m^k^*) is a normalization factor, and *m* > 1 to ensure that the series converges when *k* approaches infinity [15]. The parameter *m* controls the diffusion intensity. The smaller the *m*, the higher the degree of diffusion (default as *m* = 1.5). Notably, the operator *f_m_* has described a signal enhancement process by accumulating the transition probabilities of all lengths of paths between the two vertexes, which can also be regarded as a process of signal diffusion.

Then we defined two auxiliary operators *g* and *h* to help us map the diffusion process of the random walk defined by the operator *f_m_* to the diffusion process on the graph [15]. Denote *D_w_* as the diagonal degree matrix of the input network *W*, then *g*(*W*) defines a random walk on the graph whose (weighted) adjacency matrix is *W*:

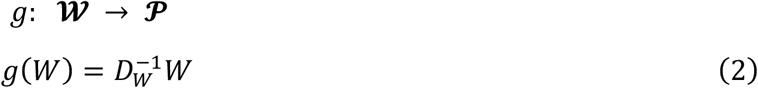

The operator *h* has the opposite effect of *g*, which recovers the underlying graph of the random walk defined by the transition matrix *P*:

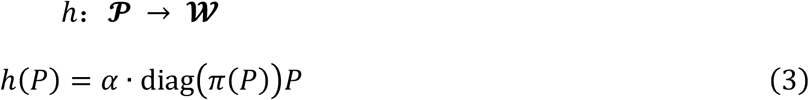

where *π*(*P*) = (*π*_1_, …, *π_n_*) is the stationary distribution of transition matrix *P* such that *πP* = *π*. When *P* defines an irreducible and aperiodic random walk, *π*(*P*) exists and can be guaranteed to be unique (that is generally true in practical applications). diag(*x*) means the diagonal matrix whose diagonal element is the vector *x*, and *α* is a constant which controls the sum of weight matrix *h*(*P*).The operator *h* multiplies the transition probability *P_ij_* by the stationary distribution of node *i* which reflects the degree information of the graph.

Formally, to map the diffusion process of random walk on the graph defined by *f_m_* onto the diffusion process on the graph, NR wraps the operator *f_m_* in operators *g* and *h* to get the composite operator *F_m_*:

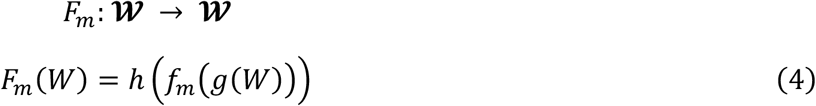

where operators *g* and *h* realize the conversion between the graph and random walk on the graph, and *f_m_* realize the diffusion of random walk on the graph.

### The difference between NR and Katz Index

We have proven in previous work that the graph operator of NR can be further written as [15]:

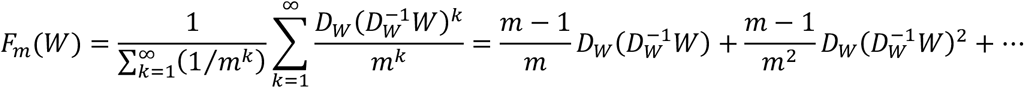

And the Katz index was calculated by [9]:

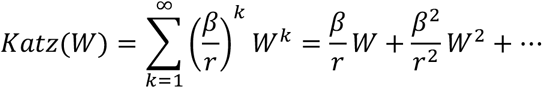

The difference between the two diffusion methods can be understood from two perspectives. First, as we described in our previous work [15], NR is a degree-normalized version of the Katz index, which makes the path with higher intermediate node degrees less important because their information is dispersed through more adjacent edges. Second, we can compute NR by computing the Katz index with a few more processing steps. When *W* is an arbitrary matrix, we cannot construct a direct connection between the two methods. But when we take a pre-processing step 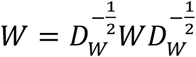, the spectral radius of the input matrix can be guaranteed to be one, which means the *r* in the formula of Katz index is one and thus have:

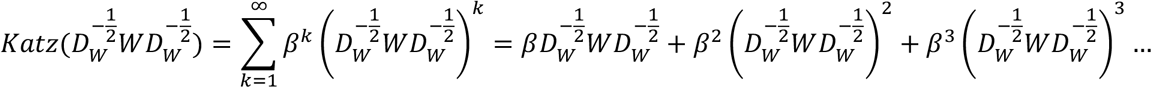

Note that the formula of NR can be further expressed as:

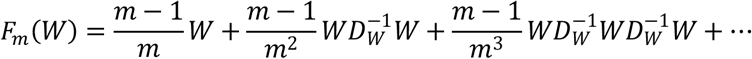

It can be seen that when 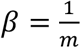, we have:

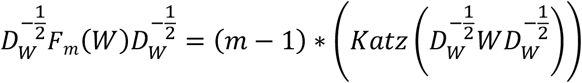

### Dimensionality reduction

We used the Uncentred Principal Component Analysis (UCPCA) [66] to obtain the low-dimensional embeddings of the corresponding cells and peaks in this study. The number of kept components was determined based on the proportion of variance that can be explained [21]. Specifically, we kept the first 50 (if peaks number less than 50,000) or 100 (if peaks number more than 50,000) principal components (PCs) and plotted the standard deviation (SD) of each PC. The kept number of PCs was determined based on the SD variance of PCs that were not selected, i.e., if the SD of the remaining PCs does not change much, we did not keep them. We give in Supplementary Fig. 29 the SD plots and the number of PCs retained on all scATAC-seq datasets in this study. Finally, we applied L2-normalization to each of the low-dimensional representations, which was a classical processing step [6][21][22] after dimensionality reduction.

### Clustering and visualization

We used the ‘scanpy.tl.louvain’ function from the Scanpy [67] python package to implement the Louvain algorithm for clustering cells, and the ‘scanpy.pl.tsne’ and ‘scanpy.pl.umap’ functions of Scanpy python package to visualize cells in two-dimensional coordinates, using the t-SNE basis and UMAP basis.

### Evaluation metrics

We used the Normalized Mutual Information (NMI) and the Adjusted Rand Index (ARI) to evaluate the clustering performance. Assuming that there are two partitions *X* = {*X*, … *X_r_*} and *Y* = {*Y*_1_, … *Y_s_*} for *N* vertexes, ARI is computed as follows:

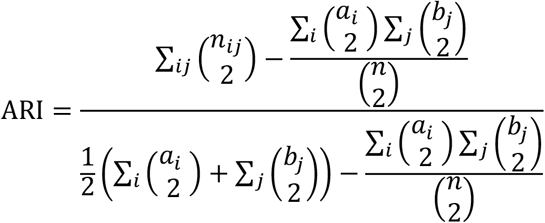

where *n_ij_* is the number of vertexes in partitions *X_i_* and *Y_i_*, and *a_i_* is the number of vertexes in partition *X_i_*, *b_j_* is the number of vertexes in partition *Y_j_*. ARI takes values from −1 to 1, and a higher value of ARI indicates better performance. Besides, NMI is computed as:

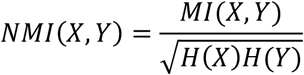

where the mutual information of two partitions and the entropy of a partition are computed as:

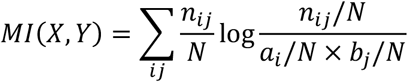

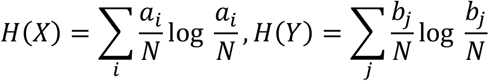

NMI takes values from 0 to 1, and a higher value of NMI indicates better performance.

We also used the silhouette coefficient to evaluate the distance between clusters, given a partition for cells. Specifically, for cell *x*:

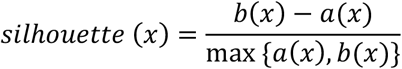

where *a*(*x*) is the average distance between *x* and other cells in the same cluster, and *b*(*x*) is the average distance between *x* and cells in the closest different cluster [65]. The distance is measured by their Euclidean distance on a two-dimensional visualization (such as UMAP and t-SNE). The silhouette coefficient of the entire partition was the average over all cells. Silhouette coefficient takes values from −1 to 1, and a higher value indicates better performance.

### Peaks annotation

Peaks in the scATAC-seq datasets were annotated using the R package ‘ChIPseeker’ [68]. In this study, the upstream and downstream regions of the gene Transcription Start Site (TSS) were defined as TSS ± 3,000 base pairs.

### Differential genes in the SOX10KD datasets

After annotating the region type of peaks, we screened 14,266 peaks (promoters of genes) in total, which were referred as background genes. So, when we talk about background genes, we are actually talking about the peaks annotated as their promoter regions. The top 50 genes with the highest correlation to *SOX10* were selected to validate the regulatory relationships with it, and we refer to them as differential genes since their accessibility changes over time. Specifically, we calculated the correlations between the *SOX10* promoter (3’UTR) and all other background genes using their low-dimensional features, and then selected the highest 50 from them. That’s to say, those differential genes may have the same (indicating positive regulatory relationship) or opposite (indicating negative regulatory relationship) co-accessibility with *SOX10*.

### Validation of cis-regulatory between genes using Cytoscape

The application Reactome [69] in the software Cytoscape [70] was used to validate the gene interactions. Specifically, we input the differential genes set, and then use the database to validate direct or indirect (by linker genes) regulatory relationships between them.

### Functional enrichment analysis

In the SOX10KD experiment, given a gene set, we used the R package ‘clusterProfiler’ [71][72] for gene functional annotation, including GO enrichment analysis (molecular function, cellular components, and biological process) and KEGG pathway enrichment analysis. Specifically, we treated the genes in the GRN obtained by Cytoscape (including those differential genes and linker genes) as foreground genes set, the background genes (14,266 genes) and linker genes as the background genes set, using the ‘enrichGO’ and ‘enrichKEGG’ functions to perform gene enrichment analysis.

### Survival analysis

Survival analysis was performed with R package ‘survival’, and the external gene expression data as well as clinical data were downloaded using the online tool Xena [73].

### Differential expression (accessibility) analysis in PBMC dataset

We used scanpy.tl.rank_genes_groups function with default settings from python package ‘Scanpy’ [67] for the standard differential expression analysis as well as differential accessibility analysis of cell subgroups in PBMC dataset.

### Gene set variation analysis (GSVA)

We built our SCARP gene signatures using the top 50 marker genes for cell subgroup 2, as shown in Fig. 3b. We used the R package ‘GSVA’ to implement the Gene set variation analysis (GSVA) with default settings for the calculation of the SCARP signature score.

### Validation of cis-regulatory using PCHi-C

PCHi-C data depicted physical interactions between the promoters (baits) and the entire genome (the other end). Since PCHi-C data were cell type-specific, we only selected those cells annotated as CD4 naive cells in the 10X Multiome data, so that there were overlapping cell types for both data types, and used SCARP to get peak embeddings. After selecting those peaks corresponding to promoters of specific genes, we got promoter-peak pairs with confidence scores calculated by cos similarity of their low-dimensional embeddings. A promoter-peak pair was considered validated by PCHi-C evidence if (1) the overlap area between promoter (from 10X Multiome) and bait (from PCHi-C) reaches half the length of one of them, or the distance between the centers of the two is less than 5kb, and (2) the overlap area between peak (from 10X Multiome) and other end (from PCHi-C) reaches half the length of one of them, or the distance between the centers of the two is less than 5kb [52]. We can then treat whether promoter-peak pairs were validated by PCHi-C as a binary classification problem and measure its accuracy by calculating AUROC score.

We used t-test to verify whether there was a significant difference in SCARP-derived cis-regulatory scores between PCHi-C validated and unvalidated promoter-peak pairs.

### Validation of cis-regulatory using ChIP-Seq

TF ChIP–seq data downloaded from ENCODE [74] was also used as another external evidence to validate cis-regulatory relationships. We took the intersections of all known human TF, promoters of PCHi-C, and the peaks annotated as promoter regions in scATAC-seq data, so that we can verify the third one with the other two. The interactions between chromosome segments were plotted using the R package ‘Cicero’ [54]. The human GRCh38 reference genome and annotation files were downloaded from ENSEMBL [75].

## Data availability

All datasets analyzed in this study were publicly available. The detailed information for the scATAC-seq and the multi-omics datasets used in this study can be found in Supplementary Table 1. The gene expression and phenotype data of the Skin Cutaneous Melanoma (SKCM) project were downloaded from TCGA. For promoter-capture Hi-C data, it can be obtained from the original publication [52][53]. For TF ChIP–seq data, it can be downloaded from the ENCODE project (https://www.encodeproject.org/) and further processed using GLUE’s script (https://github.com/gao-lab/GLUE).

## Code availability

SCARP is available at https://github.com/Wu-Lab/SCARP.

## Acknowledgements

This work has been supported by the National Key Research and Development Program of China (No. 2020YFA0712402) and the National Natural Science Foundation of China (No. 12231018).

## Author contributions

**Jiating Yu**: Conceptualization, Methodology, Software, Investigation, Formal analysis, Writing – original draft. **Duanchen Sun:** Analysis, Writing – review & editing. **Zhichao Hou:** Writing – review & editing. **Ling-Yun Wu**: Conceptualization, Methodology, Supervision, Writing – review & editing, Funding acquisition.

## Competing interests

The authors declare no competing interests.

## Additional information

Supplementary martials are available online.

## Abbreviations

PBMC: Peripheral blood mononuclear cell
HSC: Hematopoietic Stem Cells
MPP: Multipotent Progenitor
CMP: Common Myeloid Progenitor
LMPP: Lymphoid-primed Multipotent Progenitors
MEP: Megakaryocyte-erythroid Progenitor
GMP: Granulocyte-monocyte Progenitor
pDC: Plasmacytoid Dendritic Cells
CLP: Common Lymphoid Progenitor
Mono: Monocytes

## Notes

### Competing Interest Statement

The authors have declared no competing interest.

## Reference

[1] Xiong, L. et al. SCALE method for single-cell ATAC-seq analysis via latent feature extraction. Nat. Commun. 10, 1–10 (2019).

[2] Bravo González-Blas, C. et al. cisTopic: cis-regulatory topic modeling on single-cell ATAC-seq data. Nat. Methods 16, 397–400 (2019).

[3] Buenrostro, J. D. et al. Single-cell chromatin accessibility reveals principles of regulatory variation. Nature 523, 486–490 (2015).

[4] Cusanovich, D. A. et al. Multiplex single-cell profiling of chromatin accessibility by combinatorial cellular indexing. Science 348, 910–914 (2015).

[5] Cusanovich, D. A. et al. The cis-regulatory dynamics of embryonic development at single-cell resolution. Nature 555, 538–542 (2018).

[6] Dong, K. & Zhang, S. Network diffusion for scalable embedding of massive single-cell ATAC-seq data. Sci. Bull. 66, 2271–2276 (2021).

[7] Liu, Y., Zhang, J., Wang, S., Zeng, X. & Zhang, W. Are dropout imputation methods for scRNA-seq effective for scATAC-seq data? Brief. Bioinform. 23, 1–12 (2022).

[8] Schep, A. N., Wu, B., Buenrostro, J. D. & Greenleaf, W. J. ChromVAR: Inferring transcription-factor-associated accessibility from single-cell epigenomic data. Nat. Methods 14, 975–978 (2017).

[9] Katz, L. A new status index derived from sociometric analysis. Psychometrika 18, 39–43 (1953).

[10] Speed, T. P. Genetic Map Functions. Encycl. Biostat. (2005).

[11] Altshuler, D., Daly, M. J. & Lander, E. S. Genetic mapping in human disease. Science 322, 881–888 (2008).

[12] Robinson, M. A. Linkage Disequilibrium. Encycl. Immunol. 1586–1588 (1998).

[13] Teare, M. D. & Barrett, J. H. Genetic linkage studies. Lancet 366, 1036–1044 (2005).

[14] Dekker, J., Marti-Renom, M. & Mirny, L. Exploring the three-dimensional organization of genomes: interpreting chromatin interaction data. Nat. Rev. Genet. 14, 390–403 (2013).

[15] Yu, J., Leng, J. & Wu, L.-Y. Network Refinement: A unified framework for enhancing signal or removing noise of networks. Preprint at arXiv https://arxiv.org/abs/2109.09119 (2021).

[16] Yu, J. & Wu, L. Y. Multiple Order Local Information model for link prediction in complex networks. Phys. A Stat. Mech. its Appl. 600, 127522 (2022).

[17] Eraslan, G., Simon, L. M., Mircea, M., Mueller, N. S. & Theis, F. J. Single-cell RNA-seq denoising using a deep count autoencoder. Nat. Commun. 10, 1–14 (2019).

[18] van Dijk, D. et al. Recovering Gene Interactions from Single-Cell Data Using Data Diffusion. Cell 174, 716–729.e27 (2018).

[19] Buenrostro, J. D. et al. Integrated Single-Cell Analysis Maps the Continuous Regulatory Landscape of Human Hematopoietic Differentiation. Cell 173, 1535–1548.e16 (2018).

[20] Chen, S., Lake, B. B. & Zhang, K. High-throughput sequencing of the transcriptome and chromatin accessibility in the same cell. Nat. Biotechnol. 37, 1452–1457 (2019).

[21] Stuart, T. et al. Comprehensive Integration of Single-Cell Data. Cell 177, 1888–1902.e21 (2019).

[22] Stuart, T., Srivastava, A., Madad, S., Lareau, C. A. & Satija, R. Single-cell chromatin state analysis with Signac. Nat. Methods 18, 1333–1341 (2021).

[23] PBMC from a healthy donor, single cell multiome ATAC gene expression demonstration data by Cell Ranger ARC 1.0.0. 10X Genomics https://support.10xgenomics.com/single-cell-multiome-atac-gex/datasets/1.0.0/pbmc_granulocyte_sorted_10k (2020).

[24] Hie, B., Bryson, B. & Berger, B. Efficient integration of heterogeneous single-cell transcriptomes using Scanorama. Nat. Biotechnol. 37, 685–691 (2019).

[25] Zimmermann, H. W. et al. Bidirectional transendothelial migration of monocytes across hepatic sinusoidal endothelium shapes monocyte differentiation and regulates the balance between immunity and tolerance in liver. Hepatology 63, 233–246 (2016).

[26] Wierda, R. J. et al. A role for KMT1c in monocyte to dendritic cell differentiation: Epigenetic regulation of monocyte differentiation. Hum. Immunol. 76, 431–437 (2015).

[27] https://www.genome.gov/genetics-glossary/Promoter.

[28] https://en.wikipedia.org/wiki/Three_prime_untranslated_region.

[29] Chen, J. et al. Stathmin 1 is a potential novel oncogene in melanoma. Oncogene 32, 1330–1337 (2012).

[30] Fernández, N. B. et al. ROR1 contributes to melanoma cell growth and migration by regulating N-cadherin expression via the PI3K/Akt pathway. Mol. Carcinog. 55, 1772–1785 (2016).

[31] Hojjat-Farsangi, M. et al. Inhibition of the Receptor Tyrosine Kinase ROR1 by Anti-ROR1 Monoclonal Antibodies and siRNA Induced Apoptosis of Melanoma Cells. PLoS One 8, e61167 (2013).

[32] Oi, N. et al. LTA4H regulates cell cycle and skin carcinogenesis. Carcinogenesis 38, 728–737 (2017).

[33] D’Aguanno, S., Mallone, F., Marenco, M., Del Bufalo, D. & Moramarco, A. Hypoxia-dependent drivers of melanoma progression. J. Exp. Clin. Cancer Res. 40, 159 (2021).

[34] Jing, X. et al. Role of hypoxia in cancer therapy by regulating the tumor microenvironment. Mol. Cancer 18, 1–15 (2019).

[35] Levy, C., Khaled, M. & Fisher, D. E. MITF: master regulator of melanocyte development and melanoma oncogene. Trends Mol. Med. 12, 406–414 (2006).

[36] Hartman, M. L. & Czyz, M. MITF in melanoma: Mechanisms behind its expression and activity. Cell. Mol. Life Sci. 72, 1249–1260 (2015).

[37] Hwang, H. W. et al. Distinct microRNA expression signatures are associated with melanoma subtypes and are regulated by HIF1A. Pigment Cell Melanoma Res. 27, 777–787 (2014).

[38] Lakhter, A. J., Lahm, T., Broxmeyer, H. E. & Naidu, S. R. Golgi Associated HIF1a Serves as a Reserve in Melanoma Cells. J. Cell. Biochem. 117, 853–859 (2016).

[39] Feige, E. et al. Hypoxia-induced transcriptional repression of the melanoma-associated oncogene MITF. Proc. Natl. Acad. Sci. U. S. A. 108, E924–E933 (2011).

[40] Buscà, R. et al. Hypoxia-inducible factor 1α is a new target of microphthalmia-associated transcription factor (MITF) in melanoma cells. J. Cell Biol. 170, 49–59 (2005).

[41] Slominski, A. et al. The role of melanogenesis in regulation of melanoma behavior: Melanogenesis leads to stimulation of HIF-1α expression and HIF-dependent attendant pathways. Arch. Biochem. Biophys. 563, 79–93 (2014).

[42] Slominski, A., Zbytek, B. & Slominski, R. Inhibitors of melanogenesis increase toxicity of cyclophosphamide and lymphocytes against melanoma cells. Int. J. Cancer 124, 1470–1477 (2009).

[43] Sullivan, R. J. & Flaherty, K. MAP kinase signaling and inhibition in melanoma. Oncogene 32, 2373–2379 (2012).

[44] Fecher, L. A., Amaravadi, R. K. & Flaherty, K. T. The MAPK pathway in melanoma. Curr. Opin. Oncol. 20, 183–189 (2008).

[45] Lee, D. J. et al. Peroxiredoxin-2 represses melanoma metastasis by increasing E-cadherin/β-catenin complexes in adherens junctions. Cancer Res. 73, 4744–4757 (2013).

[46] Korla, P. K. et al. Somatic mutational landscapes of adherens junctions and their functional consequences in cutaneous melanoma development. Theranostics 10, 12026–12043 (2020).

[47] Slominski, A. et al. The role of melanogenesis in regulation of melanoma behavior: Melanogenesis leads to stimulation of HIF-1α expression and HIF-dependent attendant pathways. Arch. Biochem. Biophys. 563, 79–93 (2014).

[48] Malekan, M., Ebrahimzadeh, M. A. & Sheida, F. The role of Hypoxia-Inducible Factor-1alpha and its signaling in melanoma. Biomed. Pharmacother. 141, 111873 (2021).

[49] Moss, A. L. H. & Rees, M. J. W. Metastatic malignant melanoma in muscle. Br. J. Plast. Surg. 37, 250–252 (1984).

[50] Baruthio, F., Quadroni, M., Rüegg, C. & Mariotti, A. Proteomic analysis of membrane rafts of melanoma cells identifies protein patterns characteristic of the tumor progression stage. Proteomics 8, 4733–4747 (2008).

[51] Shi, X. et al. Prognostic and immune-related value of STK17B in skin cutaneous melanoma. PLoS One 17, 1–21 (2022).

[52] Cao, Z. J. & Gao, G. Multi-omics single-cell data integration and regulatory inference with graph-linked embedding. Nat. Biotechnol. 40, 1458–1466 (2022).

[53] Javierre, B. M. et al. Lineage-Specific Genome Architecture Links Enhancers and Non-coding Disease Variants to Target Gene Promoters. Cell 167, 1369–1384.e19 (2016).

[54] Pliner, H. A. et al. Cicero Predicts cis-Regulatory DNA Interactions from Single-Cell Chromatin Accessibility Data. Mol. Cell 71, 858–871.e8 (2018).

[55] McClure, B. J. et al. Pre-B acute lymphoblastic leukaemia recurrent fusion, EP300-ZNF384, is associated with a distinct gene expression. Br. J. Cancer 118, 1000–1004 (2018).

[56] Zhang, X. Y. et al. MRD-Negative Remission Induced in EP300-ZNF384 Positive B-ALL Patients by Tandem CD19/CD22 CAR T-Cell Therapy Bridging to Allogeneic Stem Cell Transplantation. Onco. Targets. Ther. 14, 5197–5204 (2021).

[57] Kuefer, M. U. et al. cDNA Cloning, Tissue Distribution, and Chromosomal Localization of Myelodysplasia/Myeloid Leukemia Factor 2 (MLF2). Genomics 35, 392–396 (1996)

[58] Heshmati, Y. et al. The chromatin-remodeling factor CHD4 is required for maintenance of childhood acute myeloid leukemia. Haematologica 103, 1169–1181 (2018).

[59] Heshmati, Y. et al. Identification of CHD4 As a Potential Therapeutic Target of Acute Myeloid Leukemia. Blood 128, 1648 (2016).

[60] Yabumoto, K. et al. Involvement of the BCL3 gene in two patients with chronic lymphocytic leukemia. Int. J. Hematol. 59, 211–218 (1994).

[61] Mckeithan, T. W. et al. BCL3 Rearrangements and t(14;19) in Chronic Lymphocytic Leukemia and Other B-Cell Malignancies: A Molecular and Cytogenetic Study. Genes Chromosom. Cancer 20, 64–72 (1997).

[62] Weinberg, J. B. et al. Apolipoprotein E (APOE) Genotype as a Determinant of Survival in Women with Chronic Lymphocytic Leukemia. Blood 110, 3081–3081 (2007).

[63] Weinberg, J. B. et al. Apolipoprotein E genotype as a determinant of survival in chronic lymphocytic leukemia. Leukemia 22, 2184–2192 (2008).

[64] Chun, E. M. et al. Expression of the apolipoprotein C-II gene during myelomonocytic differentiation of human leukemic cells. J. Leukoc. Biol. 69, 645–650 (2001).

[65] Li, Z. et al. Chromatin-accessibility estimation from single-cell ATAC-seq data with scOpen. Nat. Commun. 12, 6386 (2021).

[66] Cadima, J. & Jolliffe, I. On relationships between uncentred and column-centred principal component analysis. Pakistan J. Stat. 25, 473–503 (2009).

[67] Wolf, F. A., Angerer, P. & Theis, F. J. SCANPY: Large-scale single-cell gene expression data analysis. Genome Biol. 19, 1–5 (2018).

[68] Yu, G., Wang, L. G. & He, Q. Y. ChIPseeker: an R/Bioconductor package for ChIP peak annotation, comparison and visualization. Bioinformatics 31, 2382–2383 (2015).

[69] Croft, D. et al. Reactome: a database of reactions, pathways and biological processes. Nucleic Acids Res. 39, D691–D697 (2011).

[70] Shannon, P. et al. Cytoscape: A Software Environment for Integrated Models of Biomolecular Interaction Networks. Genome Res. 13, 2498–2504 (2003).

[71] Yu, G., Wang, L. G., Han, Y. & He, Q. Y. ClusterProfiler: An R package for comparing biological themes among gene clusters. Omi. A J. Integr. Biol. 16, 284–287 (2012).

[72] Wu, T. et al. clusterProfiler 4.0: A universal enrichment tool for interpreting omics data. Innov. 2, 100141 (2021).

[73] https://xena.ucsc.edu/.

[74] Davis, C. A. et al. The Encyclopedia of DNA elements (ENCODE): data portal update. Nucleic Acids Res. 46, D794–D801 (2018).

[75] https://asia.ensembl.org/index.html.

