## Supplementary Materials for "Single-Cell ATAC-seq analysis via Network Refinement with peaks location information"

### 1. Dataset descriptions

Here we give some detailed information on all the datasets used in this study. The number of cells, peaks, and cell clusters as well as links to the source studies for each dataset were shown in Supplementary Table 1.

- We have tested SCARP on nine benchmarking scATAC-seq datasets (Leukemia, GM12878vsHEK, InSilico, GM12878vsHL, Forebrain, Breast Tumor, Splenocyte, SOX10KD, blood2K) with reference cell labels. The first seven scATAC-seq datasets were downloaded from the corresponding source study, whose peaks had already been filtered. The SOX10KD and blood2K scATAC-seq datasets have a large number of peaks, so we filtered their peaks to different levels using different methods (based on count or variance) to test the robustness of SCARP. For example, "SOX10KD_filter_var0.6" means that we only kept peaks with variance greater than 0.6 quartiles of all peak variances, and "blood2K_filter60" means that we only kept peaks with the number of counts greater than 60.
- We used the SNARE-seq profiling of adult mouse cerebral cortex to verify the consistency of cell clustering results between gene expression and chromatin accessibility data. The cell type annotations were obtained by running the vignette downloaded from https://satijalab.org/signac/articles/snareseq.html.
- The PBMC single cell multiome (ATAC and Gene Expression) dataset from 10X Genomics was used to further gain insight into identifying new cell subpopulations and transcriptional regulation. We followed the processing step in https://github.com/gao-lab/GLUE to get the cell type annotations (using the R package "Seurat").
- To validate the biological significance of CD14 monocyte cell subtypes, we use two external scRNA-seq datasets: Human CD14 monocytes from 10X Genomics and time-series M-CSF stimulation CD14 monocytes with four donors on three timestamps (0, 3, 6 days).

<table>
<tr><th colspan="2">Dataset name</th><th>#Cells</th><th>#Peaks/ #Genes</th><th>Sparsity</th><th>#Clusters</th><th>Source study</th></tr>
<tr><td colspan="2">Leukemia</td><td>391</td><td>7602</td><td>5%</td><td>6</td><td rowspan="7">https://doi.org/10.1038/s41467-019-12630-7</td></tr>
<tr><td colspan="2">GM12878vsHEK</td><td>526</td><td>12938</td><td>4.3%</td><td>3</td></tr>
<tr><td colspan="2">InSilico</td><td>828</td><td>13668</td><td>17.5%</td><td>6</td></tr>
<tr><td colspan="2">GM12878vsHL</td><td>597</td><td>10431</td><td>3.5%</td><td>3</td></tr>
<tr><td colspan="2">Forebrain</td><td>2088</td><td>11285</td><td>4%</td><td>8</td></tr>
<tr><td colspan="2">Breast Tumor</td><td>384</td><td>27884</td><td>2.7%</td><td>4</td></tr>
<tr><td colspan="2">Splenocyte</td><td>3166</td><td>77294</td><td>16.6%</td><td>12</td></tr>
<tr><td colspan="2">SOX10KD</td><td>598</td><td>78659</td><td>4.6%</td><td>8</td><td>https://doi.org/10.1038/s41592-019-0367-1</td></tr>
<tr><td colspan="2">blood2K</td><td>2034</td><td>134962</td><td>5%</td><td>9</td><td>https://doi.org/10.1016/j.cell.2018.03.074</td></tr>
<tr><td colspan="2">SNARE-seq (Adult mouse cerebral)</td><td>8055</td><td>243838</td><td>1.1%</td><td>13</td><td>https://www.nature.com/articles/s41587-019-0290-0</td></tr>
<tr><td colspan="2">10X Multiome PMBC</td><td>10412</td><td>105949</td><td>7.4%</td><td>19</td><td>https://support.10xgenomics.com/single-cell-multiome-atac-gex/datasets/1.0.0/pbmc_granulocyte_sorted_10k</td></tr>
<tr><td colspan="2">10X Genomics Human CD14+ monocytes</td><td>2612</td><td>32738</td><td>1.4%</td><td>/</td><td>https://support.10xgenomics.com/single-cell-gene-expression/datasets/1.1.0/cd14_monocytes</td></tr>
<tr><td rowspan="8">M-CSF stimulation CD14+ monocytes</td><td>donor1-day0</td><td>966</td><td>12448</td><td>0.4%</td><td>/</td><td rowspan="8">https://www.ncbi.nlm.nih.gov/geo/query/acc.cgi?acc=GSE126085 https://www.nature.com/articles/s41587-019-0113-3/</td></tr>
<tr><td>donor2-day0</td><td>985</td><td>12299</td><td>0.5%</td><td>/</td></tr>
<tr><td>donor3-day0</td><td>1098</td><td>11345</td><td>0.4%</td><td>/</td></tr>
<tr><td>donor4-day0</td><td>1009</td><td>10985</td><td>0.5%</td><td>/</td></tr>
<tr><td>donor1-day3</td><td>563</td><td>17426</td><td>16.8%</td><td>/</td></tr>
<tr><td>donor2-day3</td><td>538</td><td>17145</td><td>19.4%</td><td>/</td></tr>
<tr><td>donor1-day6</td><td>1114</td><td>17887</td><td>13.6%</td><td>/</td></tr>
<tr><td>donor2-day6</td><td>955</td><td>17754</td><td>16.2%</td><td>/</td></tr>
</table>

**Supplementary Table 1 A brief description of all the datasets used in this study**. Details can be found in the corresponding source studies.

### 2. Comparable methods and their implementation details.

We compared SCARP with six other state-of-the-art methods (original, DCA, MAGIC, PCA, cisTopic, scAND). We now give some details about how we implemented them and show the selection of parameters for each method in Supplementary Table 2.

**2.1 The original method** treated peaks (cells) directly as the high-dimensional features of cells (peaks), which represented a naive approach.

**2.2 DCA (deep count autoencoder network)** is a deep learning framework designed for scRNA-seq analysis and can be easily implemented with the Scanpy python package. We followed the instructions in https://scanpy.readthedocs.io/en/stable/generated/scanpy.external.pp.dca.html#scanpy.external.pp.dca and set the parameter 'mode' to 'latent', the parameter 'optimizer' to 'RMSprop', and all other parameters to their default values.

**2.3 MAGIC (Markov affinity-based graph imputation of cells)** used diffusion to denoise the scRNA-seq count matrix and can also be implemented by the Scanpy python package. We followed the instructions in https://scanpy.readthedocs.io/en/stable/generated/scanpy.external.pp.magic.html#scanpy.external.pp.magic and set the parameter 'name_list' to 'pca_only', and all other parameters to their default values.

**2.4 cisTopic** used the Latent Dirichlet Allocation model for co-optimal clustering of cells and peaks and low-dimensional embeddings extraction. We used the R package 'cisTopic' and followed the instructions in http://htmlpreview.github.io/?https://github.com/aertslab/cisTopic/blob/master/vignettes/CompleteAnalysis.html to implement it. The number of kept components of embeddings was automatically determined.

**2.5 scAND (scATAC-seq data Analysis via Network Diffusion)** used the Katz index to fill in missing values of scATAC-seq data, and used PCA to obtain embedding of cells and peaks. We followed the instructions in http://www.zhanglab-amss.org/homepage/software/scAND_Code.rar to implement it. The parameter $\beta$ of Katz index is automatically determined, and the number of kept components is selected based on the "elbow plot". Using the eigen-decomposition technique, the computational complexity of scAND depends mainly on the calculation of top-$l$ eigen-decomposition of the adjacency matrix with $N$ nodes and $M$ edges, which is $O(T(Nl^2 + Ml))$ ($N$ is the total number of peaks and cells, $M$ is twice the number of non-zero counts of scATAC-seq data, and $T$ is the iterations times).

**2.6 PCA (Principal Component Analysis)** was implemented by the sklearn python package. The number of kept components was selected in the same way as for scAND.

| Dataset name | PCA | cisTopic | scAND | | SCARP |
|---|---|---|---|---|---|
| | # Kept component | # Kept component | beta | # Kept component | # Kept component |
| **Leukemia** | 10 | 9 | 0.9 | 10 | 25 |
| **GM12878vsHEK** | 10 | 6 | 0.95 | 10 | 5 |
| **InSilico** | 20 | 8 | 0.85 | 20 | 10 |
| **GM12878vsHL** | 10 | 14 | 0.95 | 10 | 5 |
| **Forebrain** | 10 | 15 | 0.95 | 10 | 10 |
| **Breast Tumor** | 10 | 14 | 0.95 | 10 | 5 |
| **Splenocyte** | 20 | 6 | 0.75 | 20 | 20 |
| **blood2K** | 20 | 6 | 0.9 | 20 | 50 |
| **SOX10KD** | 20 | 7 | 0.9 | 20 | 25 |

**Supplementary Table 2 Parameter selection for different methods on different datasets**. For SCARP, we used the default values for each parameter in all experiments ($m = 1.5, \gamma = 3000,\ and\ \beta = 5000$).

### 3. Running time of various methods.

Here we briefly analyzed the computational complexity of SCARP and showed the running times of the different methods on nine benchmarking scATAC-seq datasets in Supplementary Table 3.

For scATAC-seq data with $c$ cells and $p$ peaks, the computational complexity of SCARP depends mainly on the computation of the NR diffusion process. Specifically, NR was performed on each subnetwork with $c$ cells and $p_k$ peaks, which resulted in the computational complexity of $O((c+p_k)^3)$. Then the total computational complexity of running NR on all subgraphs was $O\left(\sum_{k=1}^{M}(c+p_k)^3\right)$ with $M$ subgraph, which was basically $O\left(Mc^3+\sum_{k=1}^{M}p_k^3\right)$. There are a few tricks to reduce the time cost of SCARP. Firstly, NR diffusion on $M$ subgraph can be computed in parallel, which makes the NR calculation time only depend on the time used for the largest subnetwork. Second, users can filter some peaks based on counts or variance, since SCARP is robust against peaks selection, as shown in the main study. For scATAC-seq data with a large number of cells, SCARP takes more time.

| Dataset name | DCA | MAGIC | cisTopic | scAND | SCARP | SCARP+PCA |
|---|---|---|---|---|---|---|
| **Leukemia** | 22 s | 12 s | 38 s | 5 s | 3 s | 5 s |
| **GM12878vsHEK** | 46 s | 1 s | 59 s | 11 s | 7 s | 13 s |
| **GM12878vsHL** | 39 s | 11 s | 48 s | 9 s | 6 s | 15 s |
| **Breast Tumor** | 1 min 33 s | 15 s | 1 min 28 s | 16 s | 17 s | 27 s |
| **InSilico** | 2 min | 1 s | 3 min 24 s | 29 s | 10 s | 21 s |
| **Forebrain** | 2 min 33 s | 4 s | 2 min 6 s | 27 s | 23 s | 1 min 5 s |
| **SOX10KD** | 4 min 40 s | 8 s | 5 min 20 s | 1min 2 s | 2 min 20 s | 3 min |
| **blood2K** | 47 min 34 s | 1 min 18 s | 23 min 28 s | 4 min 20 s | 12 min 18 s | 19 min 10 s |
| **Splenocyte** | 45 min 40 s | 1 min 20 s | 1 h 50 min | 10 min 45 s | 3 min 20 s | 6 min 32 s |

**Supplementary Table 3 Running time of various methods**. Codes were running in Intel(R) Xeon(R) CPU E5-2640 v4 @ 2.40GHz 2.40 GHz.

### 4. Supporting results for main Fig.2

In this section, we provided some complementary results of applying SCARP on the benchmarking scATAC-seq datasets, as a supplement to Fig. 2 in the main text.

Supplementary Fig. 1 and Supplementary Fig. 2 showed the confusion matrixes of applying seven methods on nine benchmarking datasets, which displayed the consistency between the results obtained from clustering using the Louvain algorithm based on low-dimensional representations of the cells and the annotated cell labels.

Supplementary Fig. 3 and Supplementary Fig. 4 showed the UMAP plots of applying seven methods on nine benchmarking datasets. Supplementary Fig. 5 and Supplementary Fig. 6 showed the t-SNE plots of applying seven methods on nine benchmarking datasets. Cells were colored by the annotated cell types in the corresponding dataset.

Supplementary Fig. 7 showed the cell clustering performance of various methods under NMI evaluation, as a supplement to Fig. 2e in the main text.

Supplementary Fig. 9, Supplementary Fig. 10, and Supplementary Fig. 11 showed the sensitivity of SCARP's parameters $m$, $\beta$, and $\gamma$ when taking different values, respectively, which demonstrated the robustness of the SCARP to parameter selections.

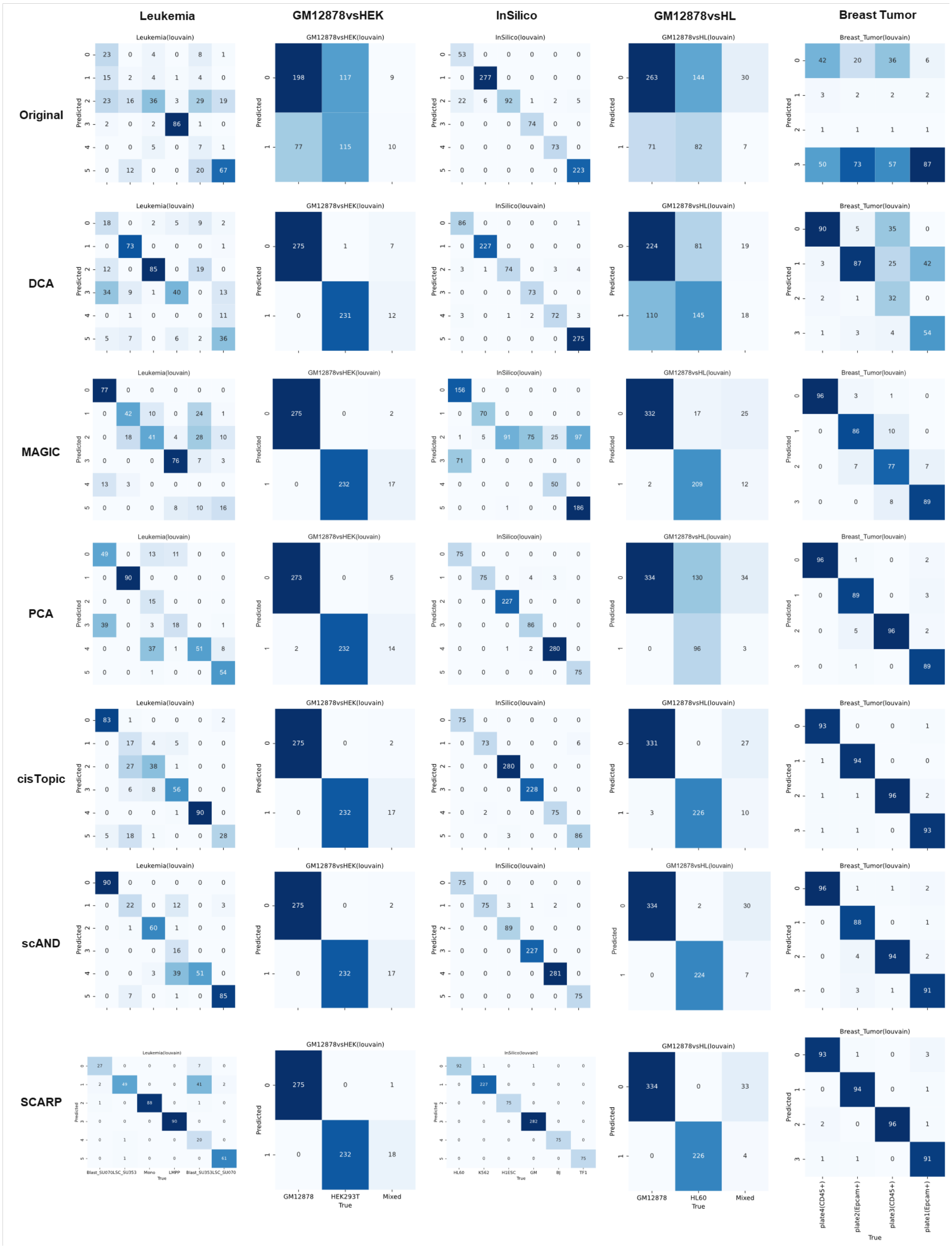

**Supplementary Fig. 1 Confusion matrixes of applying various methods on the Leukemia, GM12878vsHEK, InSilico, GM12878vsHL and Breast Tumor scATAC-seq datasets.**

The legends are the same as in Fig. 2c of the main text.

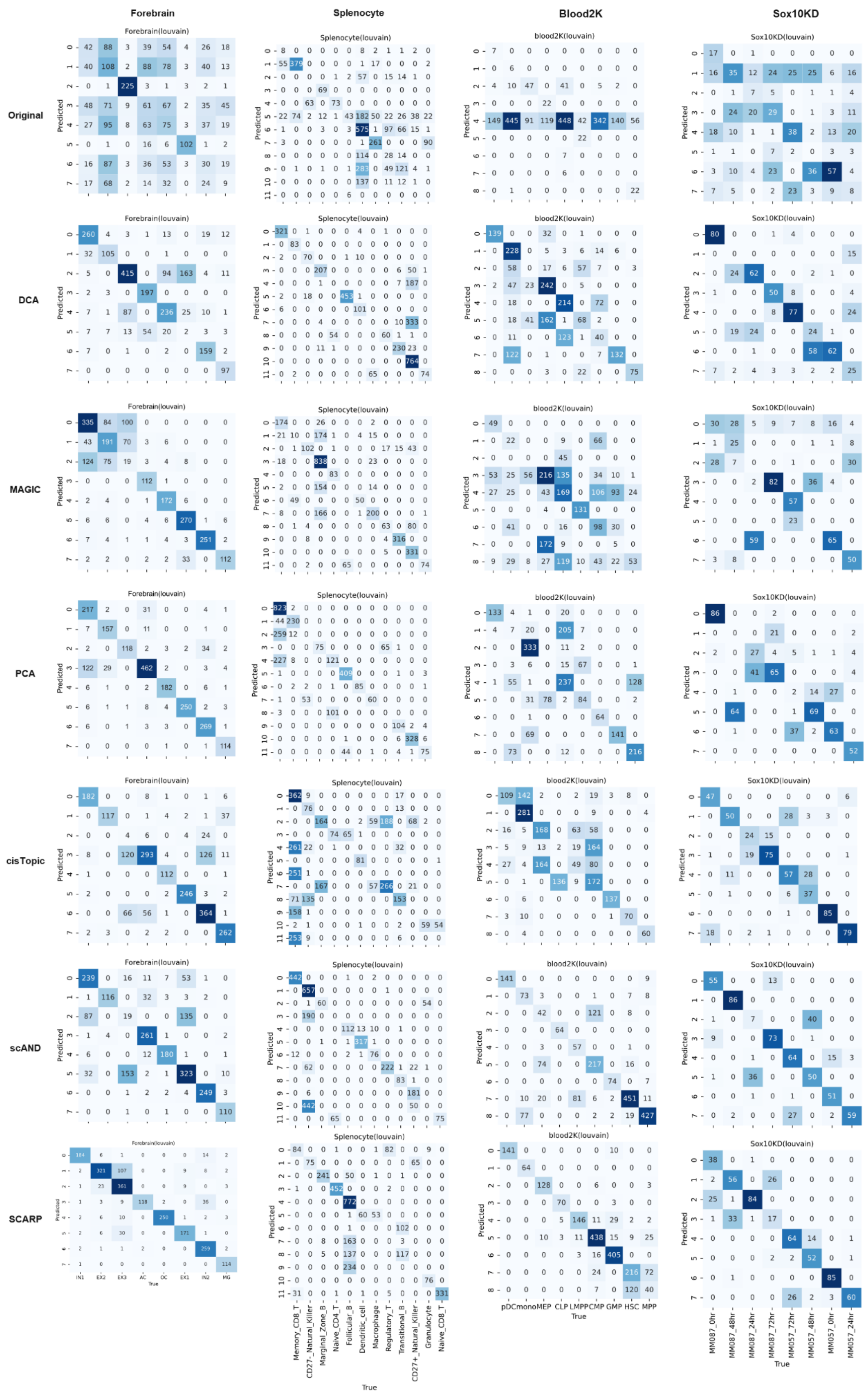

**Supplementary Fig. 2 Confusion matrixes of applying various methods on the Forebrain, Splenocyte, Blood2K, and SOX10KD scATAC-seq datasets.**

The legends are the same as in Fig. 2c of the main text.

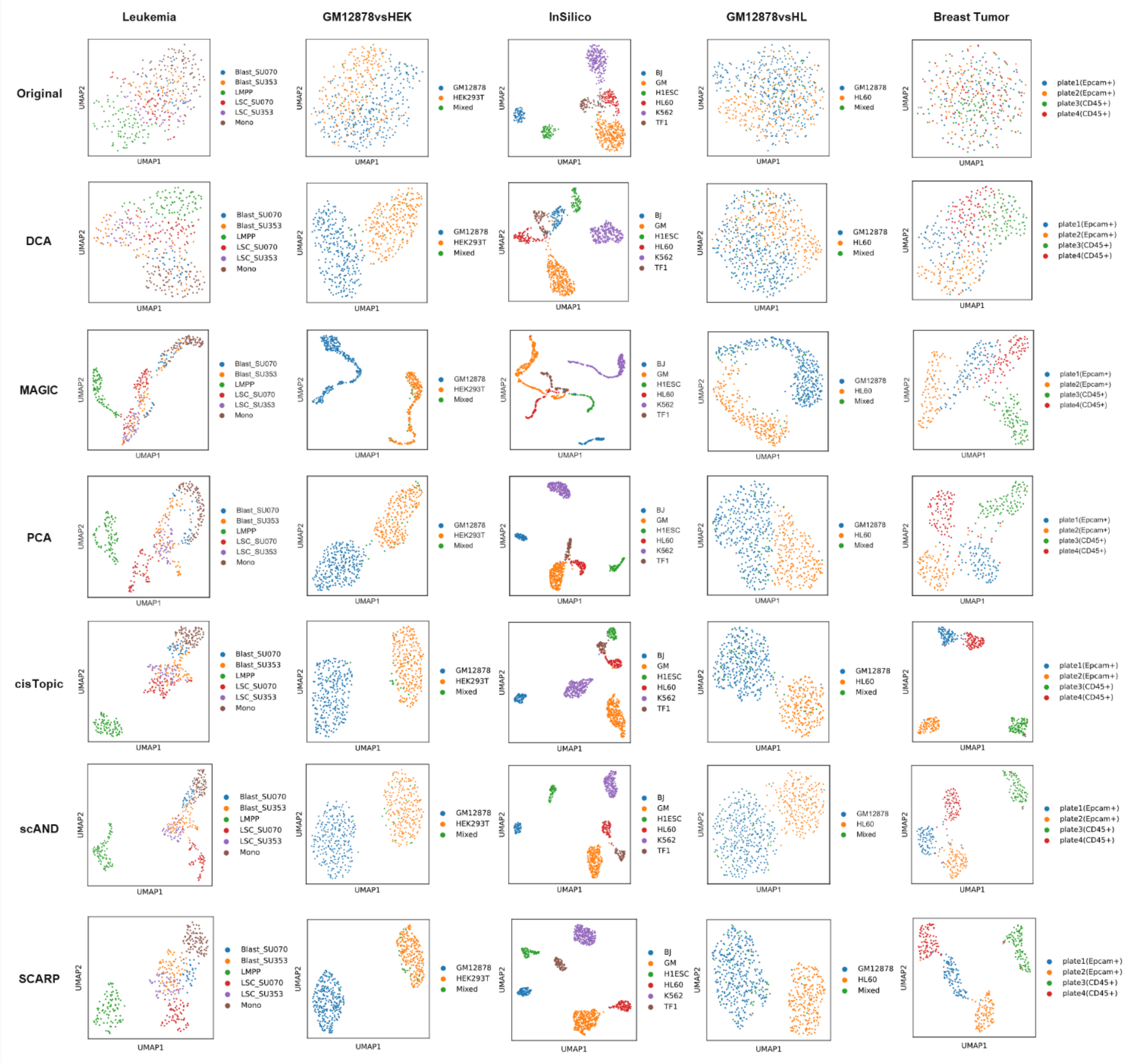

**Supplementary Fig. 3 UMAP plots of applying various methods on the Leukemia, GM12878vsHEK, InSilico, GM12878vsHL, and Breast Tumor scATAC-seq datasets.**

The cells were colored by the annotated cell types in corresponding dataset.

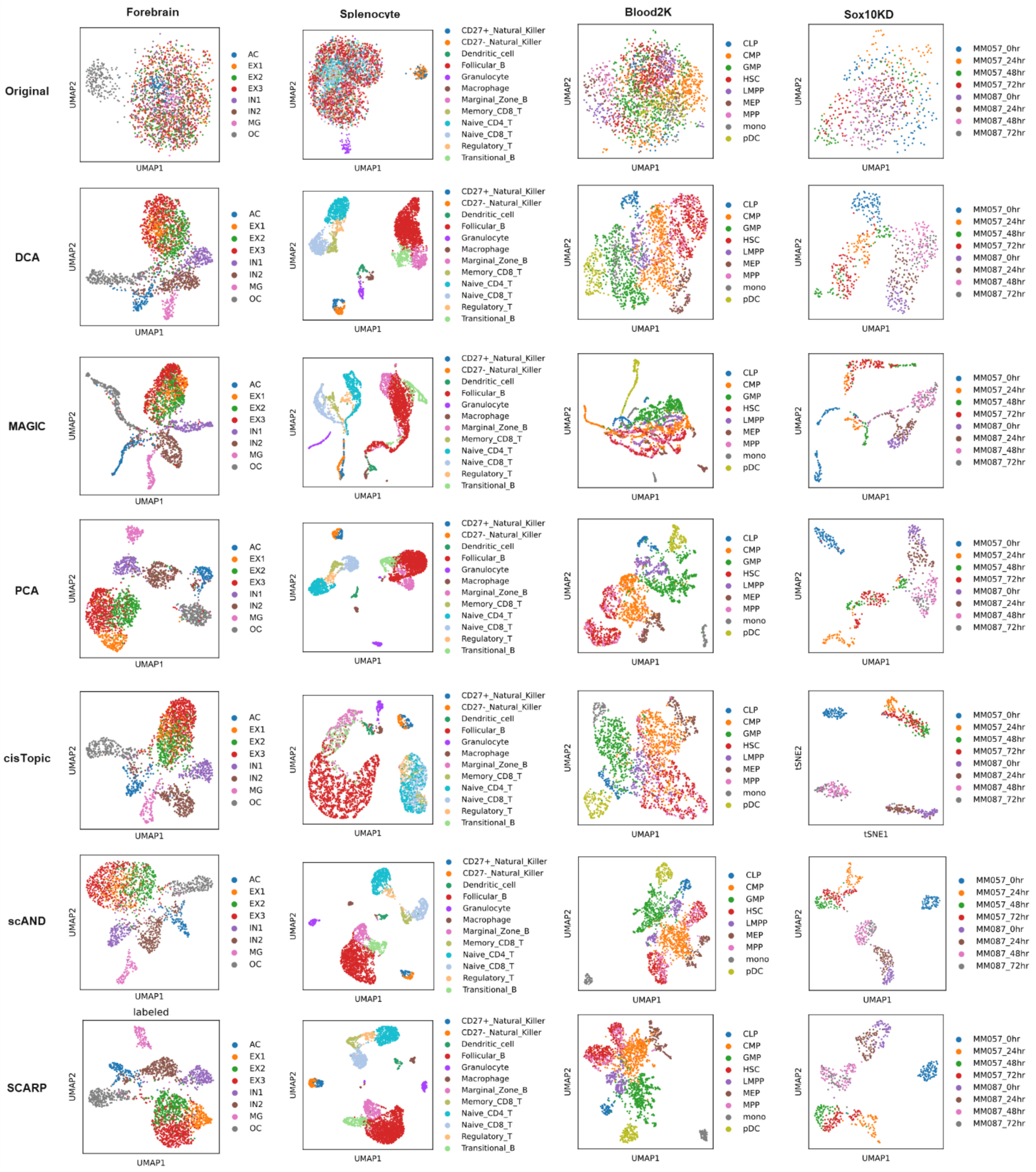

**Supplementary Fig. 4 UMAP plots of applying various methods on the Forebrain, Splenocyte, Blood2K, and SOX10KD scATAC-seq datasets.**

The cells were colored by the annotated cell types in corresponding dataset.

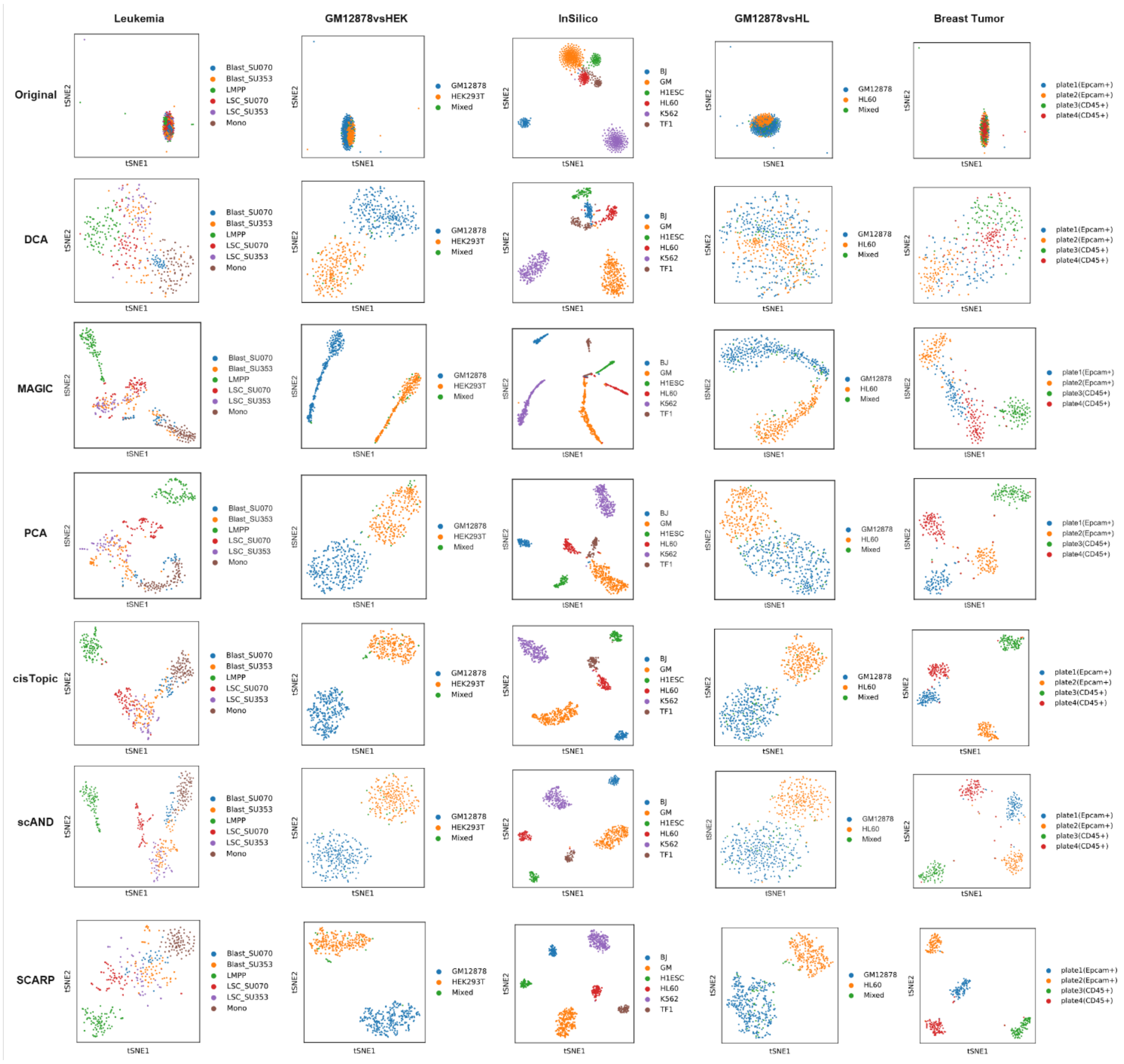

**Supplementary Fig. 5 TSNE plots of applying various methods on the Leukemia, GM12878vsHEK, InSilico, GM12878vsHL, and Breast Tumor scATAC-seq datasets.**

The cells were colored by the annotated cell types in corresponding dataset.

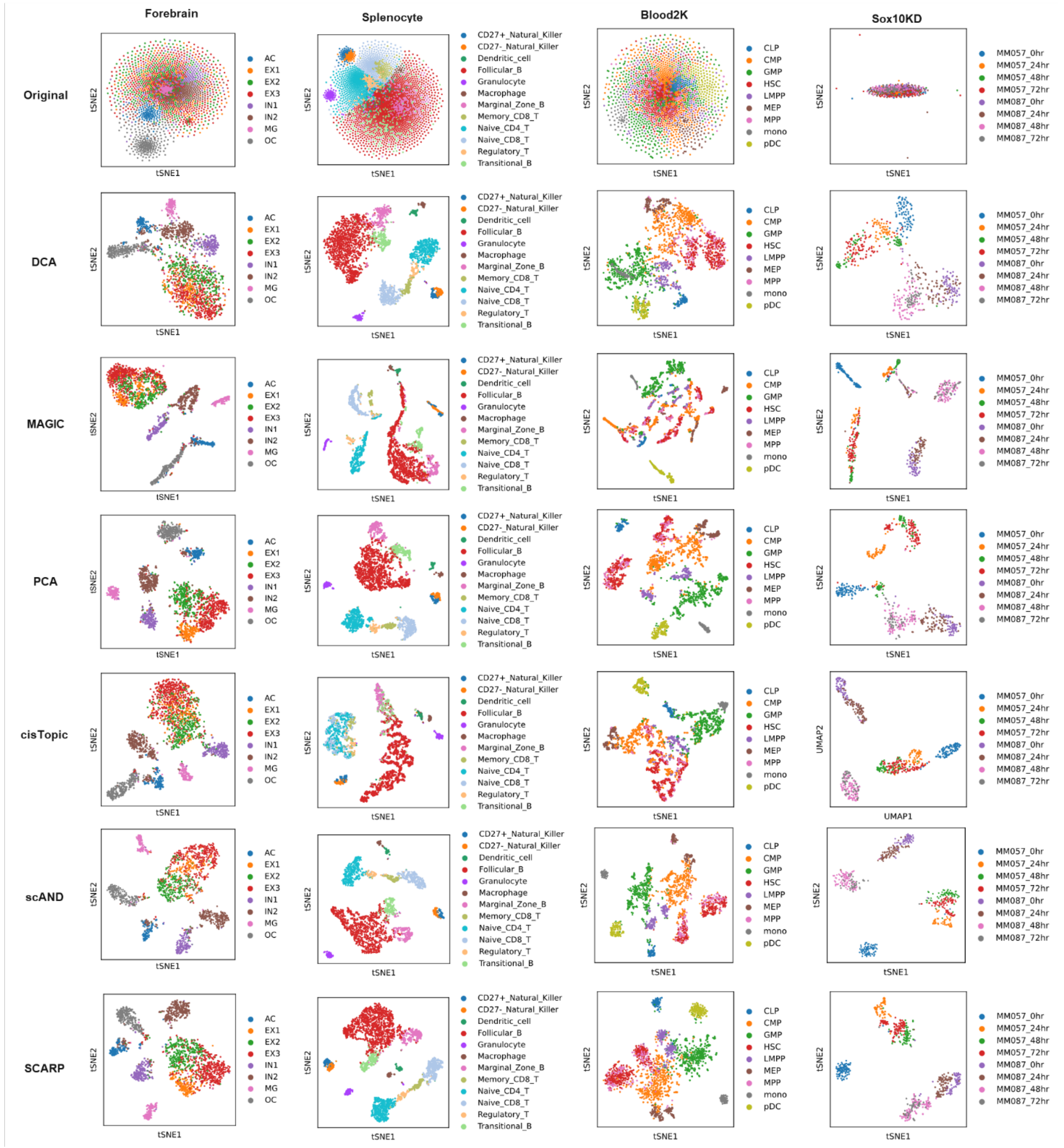

**Supplementary Fig. 6 TSNE plots of applying various methods on the Forebrain, Splenocyte, Blood2K, and SOX10KD scATAC-seq datasets.**

The cells were colored by the annotated cell types in corresponding dataset.

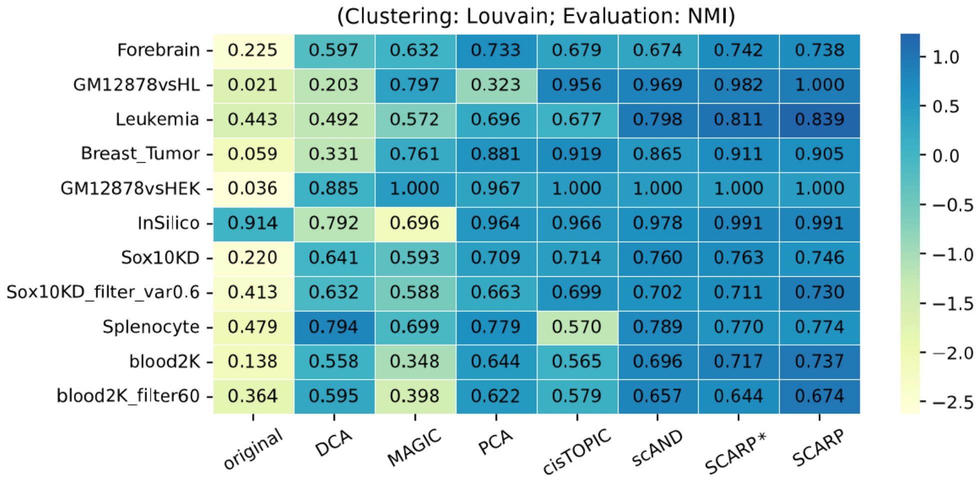

**Supplementary Fig. 7 The cell clustering performance of applying eight methods (x-axis) on 11 scATAC-seq datasets (y-axis) under NMI evaluation, as a supplement to the Fig. 2e in the main text.**

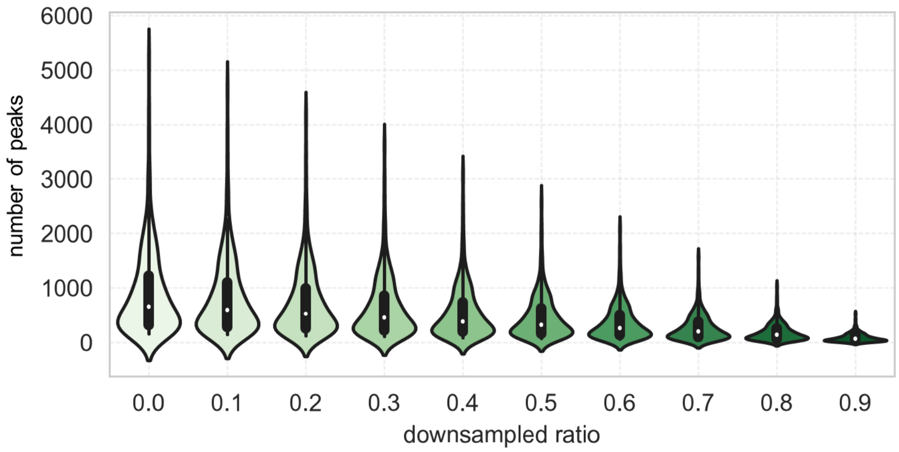

**Supplementary Fig. 8 The number of peaks (y-axis) at different downsampling ratios (x-axis) on the Leukemia scATAC-seq dataset, which led to different number of peaks.**

Specifically, by downsampling the Leukemia scATAC-seq dataset, some count values that were original non-zero become zero, resulting in some peaks being inaccessible on all cells, and we removed those peaks to obtain the new dataset.

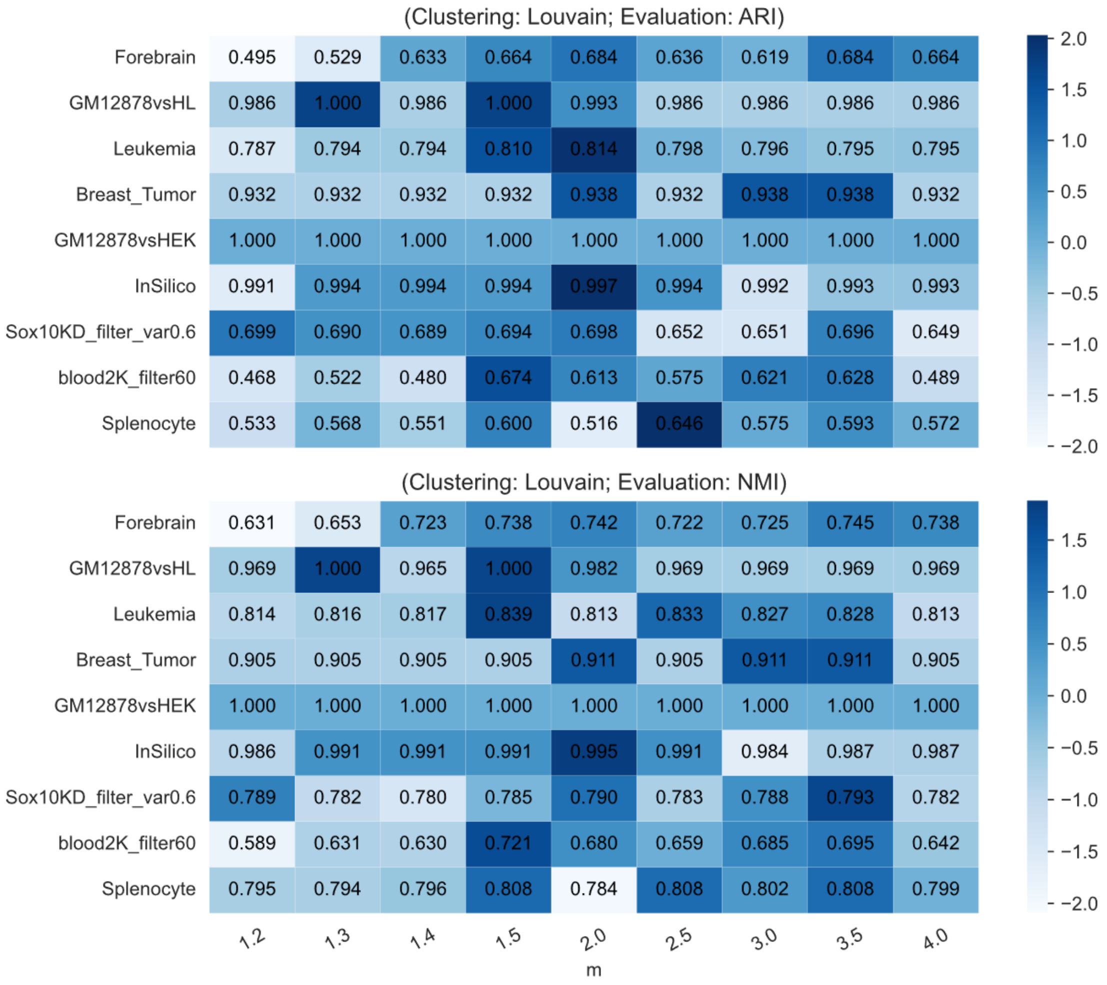

**Supplementary Fig. 9 Sensitivity of parameter $m$ of SCARP when taking different values (x-axis) on various scATAC-seq datasets (y-axis), evaluated by ARI (upper panel) and NMI (lower panel) evaluations.**

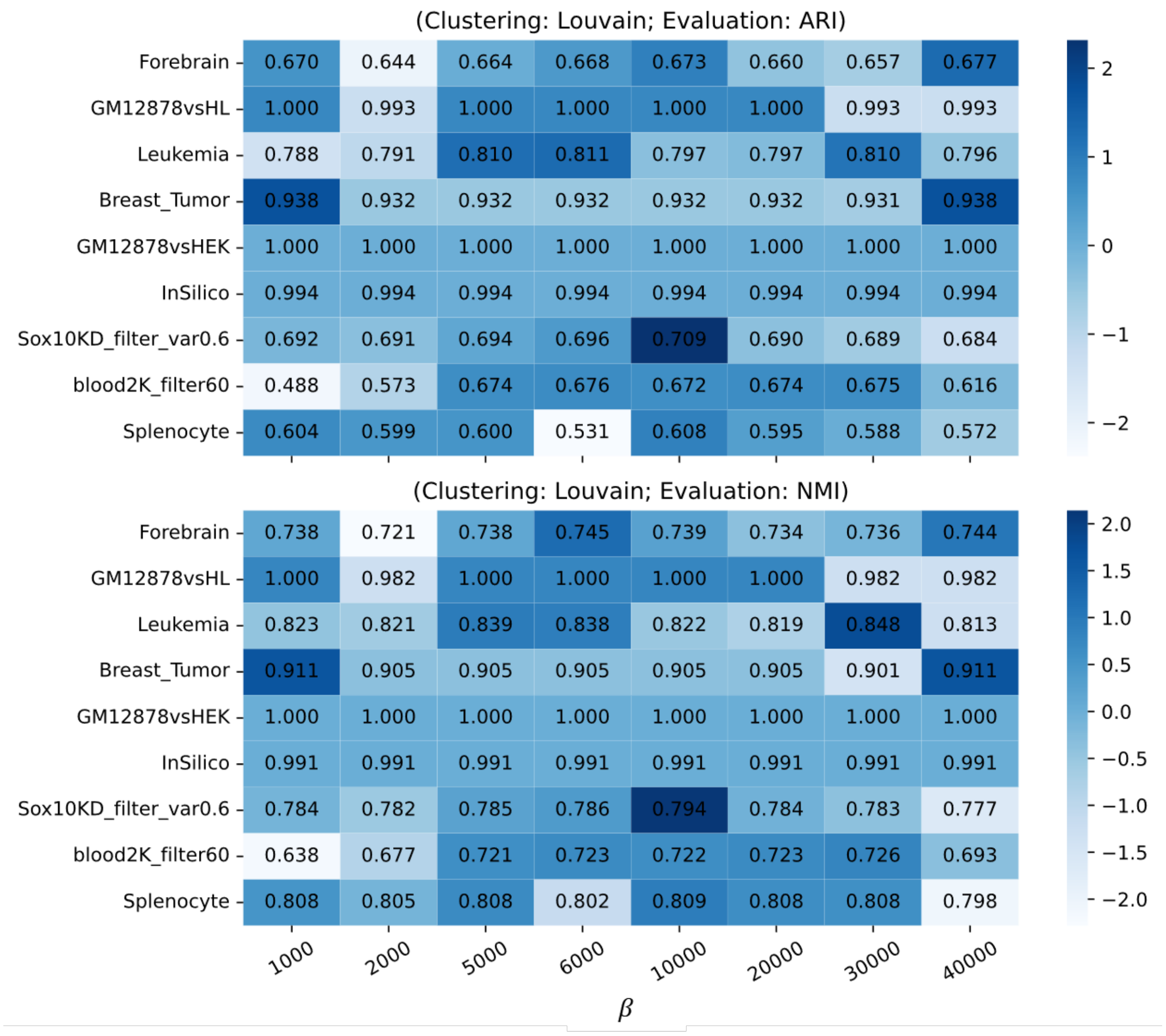

**Supplementary Fig. 10 Sensitivity of parameter $\beta$ of SCARP when taking different values (x-axis) on various scATAC-seq datasets (y-axis), evaluated by ARI (upper panel) and NMI (lower panel) evaluations.**

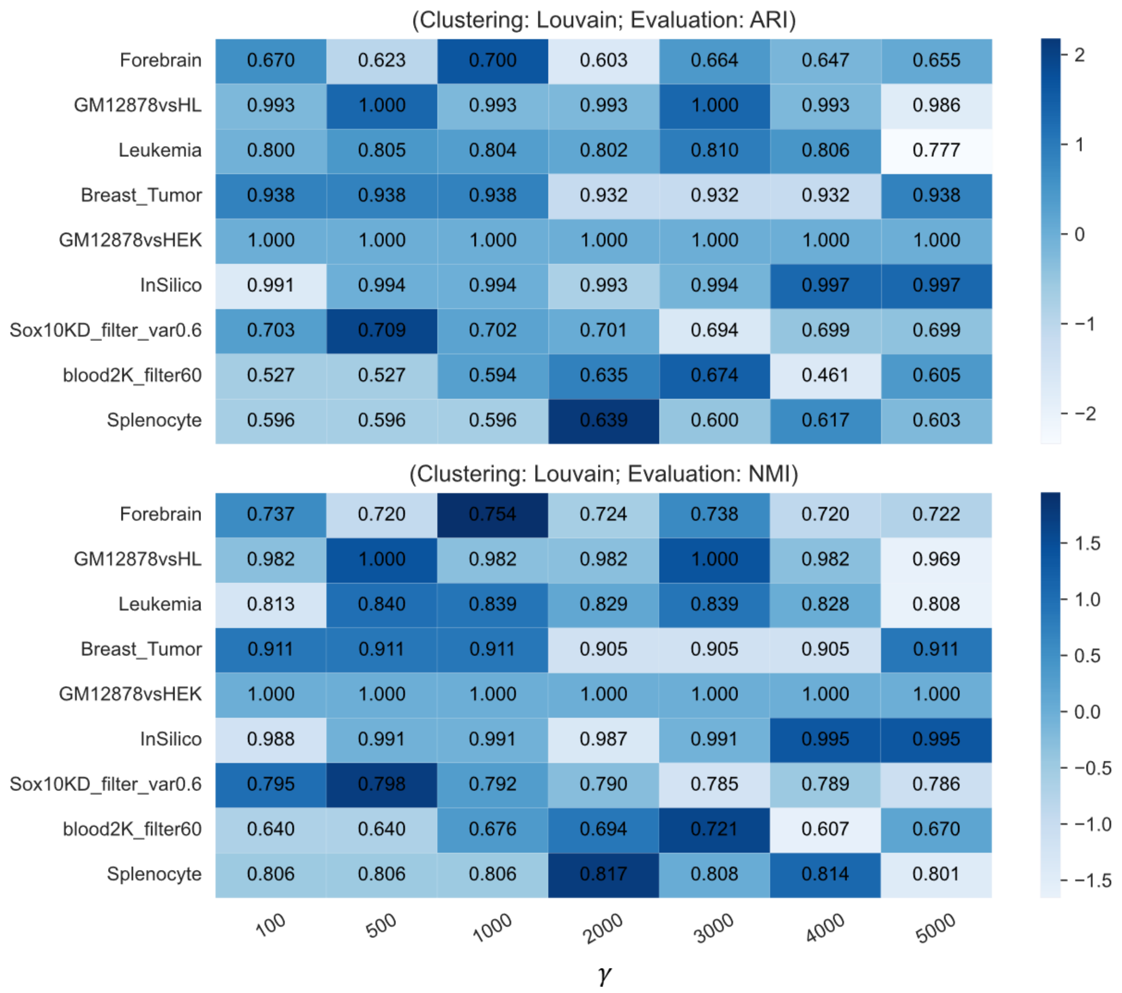

**Supplementary Fig. 11 Sensitivity of parameter $\gamma$ of SCARP when taking different values (x-axis) on various scATAC-seq datasets (y-axis), evaluated by ARI (upper panel) and NMI (lower panel) evaluations.**

### 5. Supporting results for main Fig.3

In this section, we give some supplementary results for how SCARP reveals new biologically significant cell subpopulations, and how we validated this new cell subtype using external CD14 monocytes scRNA-seq data.

We first used Seurat to process the 10X Multiome Peripheral blood mononuclear cell (PBMC) dataset and obtain cell type annotations, as shown in Supplementary Fig. 12. Yet we found that the cells originally annotated as CD14 Monocytes were clearly divided into two groups under the UMAP visualization of SCARP. We then performed a standard gene differential expression analysis as well as peak differential accessibility analysis to find marker genes as well as marker peaks of each subgroup. Supplementary Fig. 13 shows some of the marker genes and marker peaks for two cell clusters, and Supplementary Fig. 14 visualizes the expression of some marker genes and marker peaks to further show the biological differences between two cell clusters.

We further used two external CD14 monocyte scRNA-seq datasets to ascertain that the two cell subgroups identified by SCARP are indeed biologically significant, one from human CD14 monocytes stimulated by macrophage colony-stimulating factor (M-CSF) at days 0, 3, and 6, which had four donors (replicates), and the other from 10X Genomics CD14 monocytes scRNA-seq dataset. Some supplement results for M-CSF CD14 monocytes RNA-seq were given in Supplementary Fig. 15 and Supplementary Fig. 16, and supplement results for 10X Genomics CD14 monocytes RNA-seq were given in Supplementary Fig. 17.

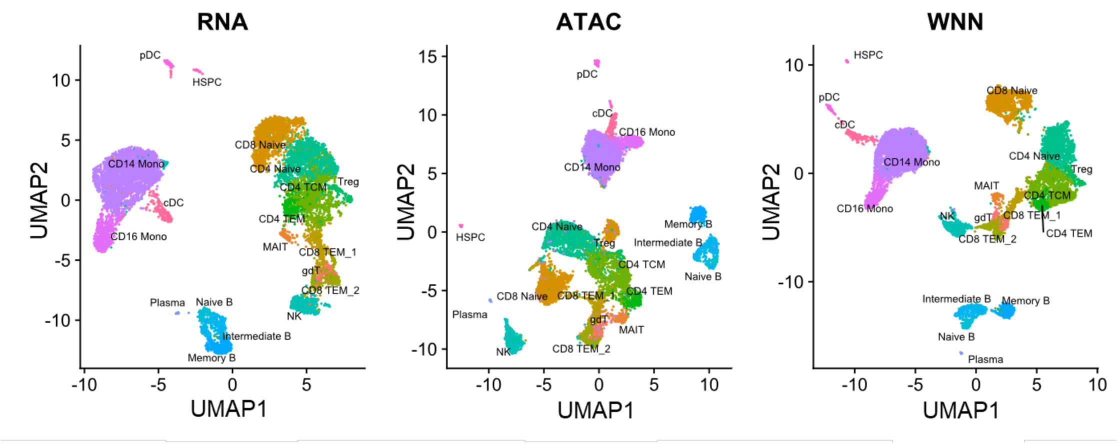

**Supplementary Fig. 12 The original cell type annotation results on 10X Multiome Peripheral blood mononuclear cell PBMC dataset.**

This is obtained by running code from [https://github.com/gao-lab/GLUE](https://github.com/gao-lab/GLUE). Obviously, all the CD14 monocytes are clustered together, and no cell subtypes are founded.

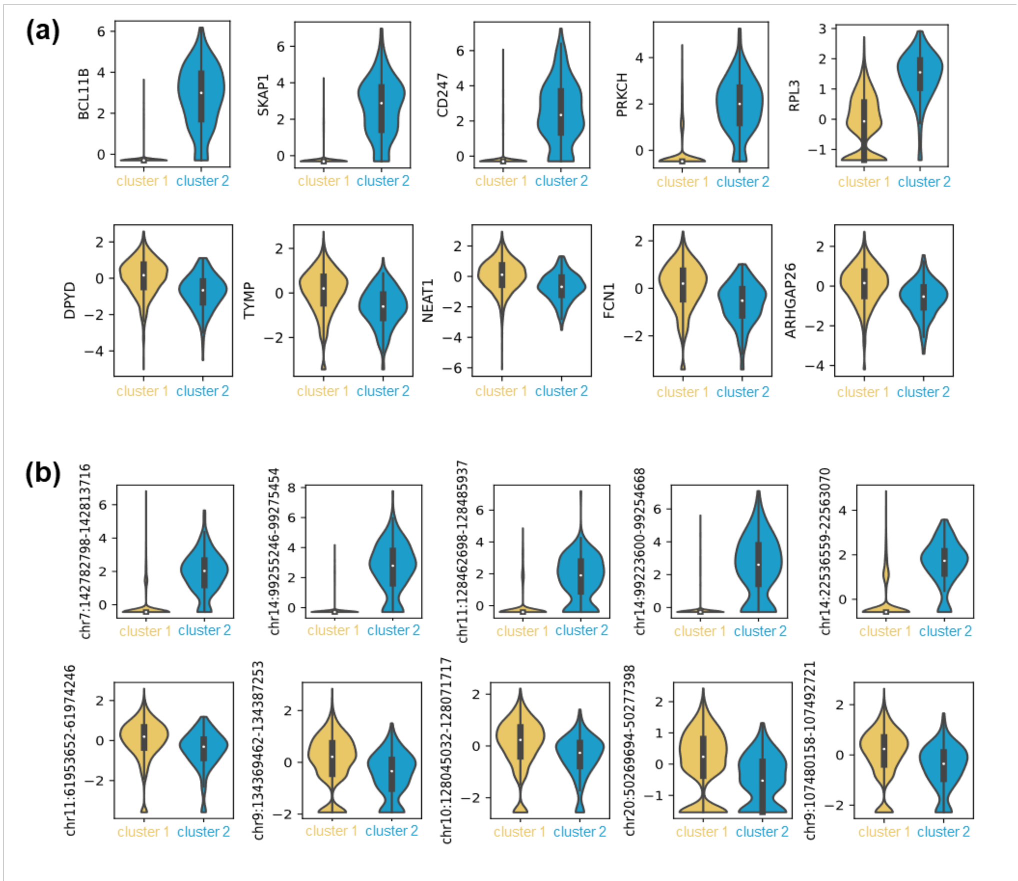

**Supplementary Fig. 13 Difference between cell subpopulations revealed by SCARP.**

**(a)** Top 5 marker genes for cell cluster 2, and top 5 marker genes for cell cluster 1. **(b)** Top 5 marker peaks for cell cluster 2, and top 5 marker peaks for cell cluster 1.

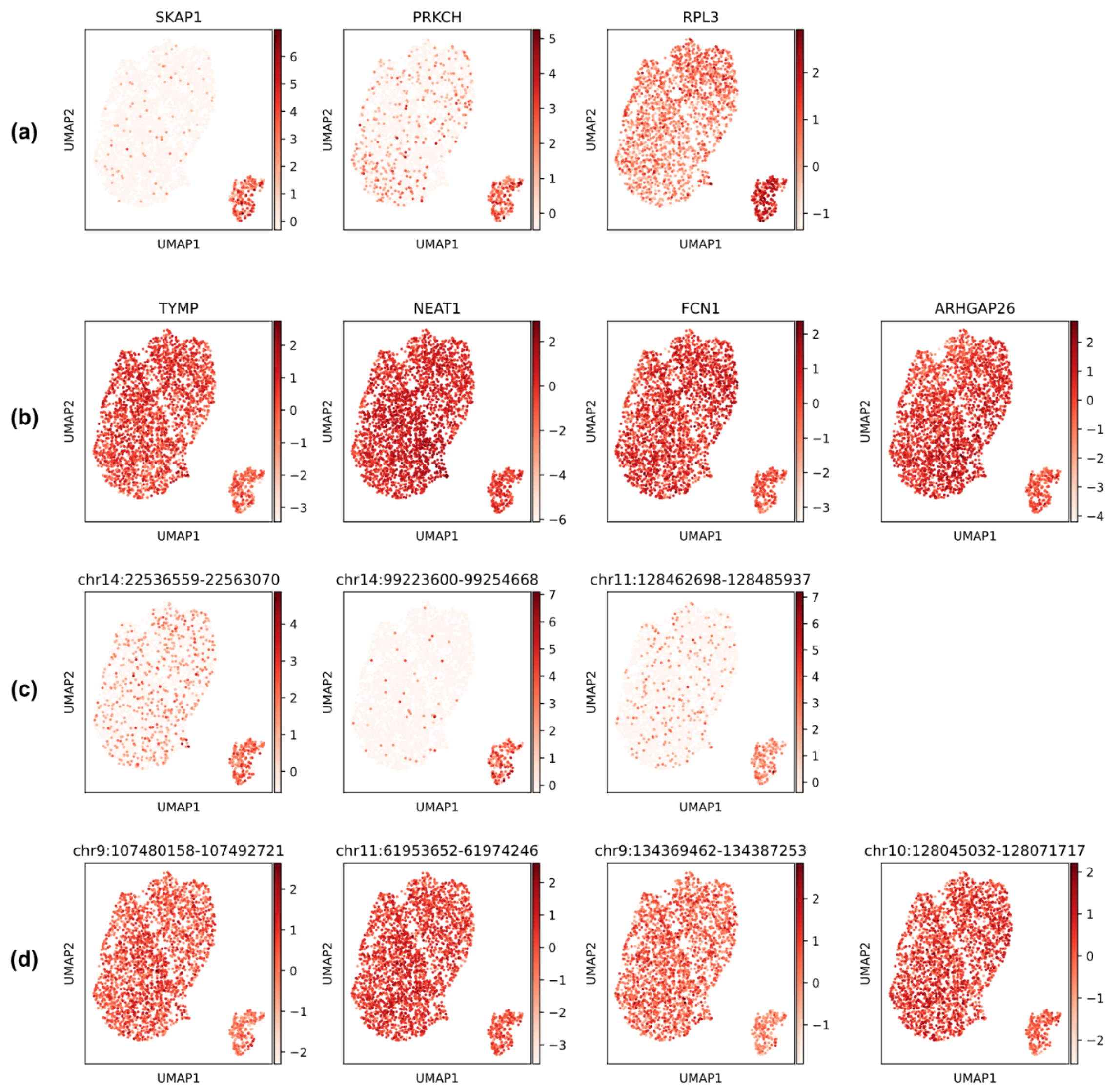

**Supplementary Fig. 14 The UMAP plots colored by the expression of marker genes and accessibility of marker peaks.**

The UMAP plots colored by the expression of **(a)** marker genes *SKAP1*, *PRKCH*, and *RPL3* of cells subgroup 2, **(b)** marker genes *TYMP*, *NEAT1*, *FCN1*, and *ARHGAP26* of cells subgroup 1, and colored by the accessibility of **(c)** marker peaks chr14: 22536559-22563070, chr14: 99223600-99254668, chr11: 128462698-128485937 for cells subgroup 2, **(d)** marker peaks chr chr9:107480158-107492721, chr11: 61953652-61974246, chr9: 134369462-134387253, and chr10: 128045032-128071717 for cells subgroup 1.

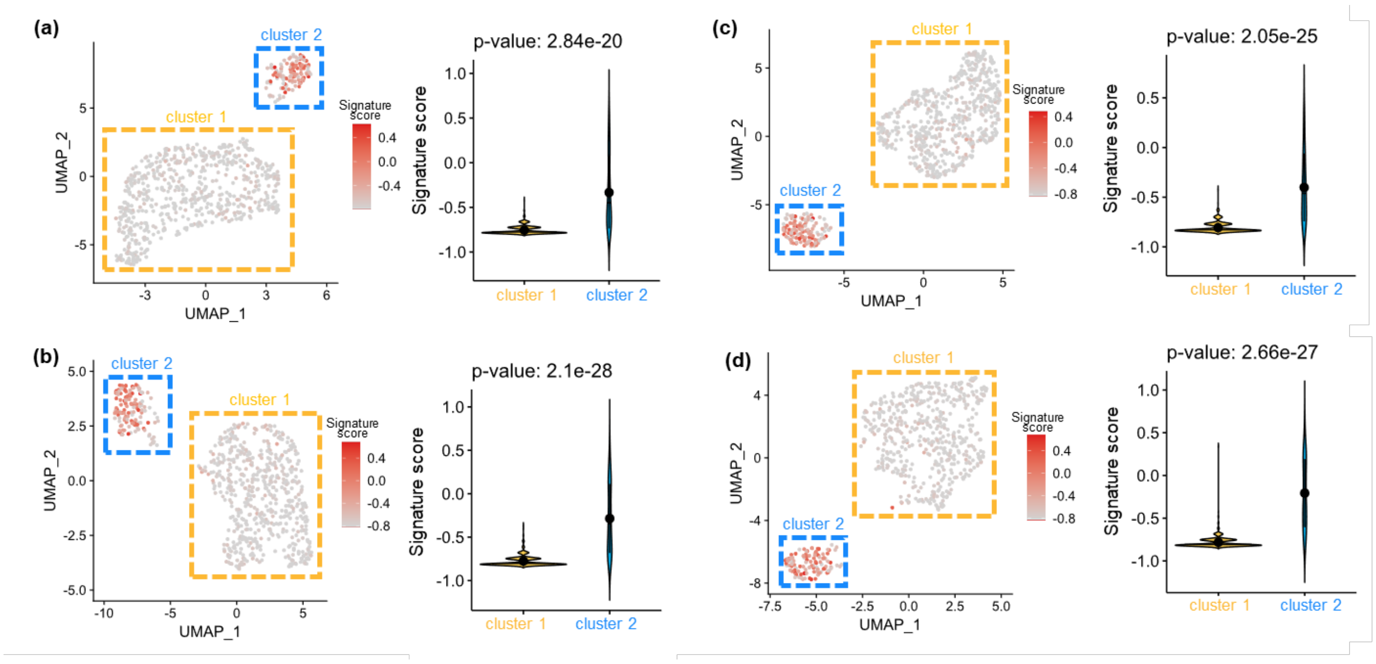

**Supplementary Fig. 15 Results on the M-CSF stimulating CD14 monocytes scRNA-seq dataset, four replicates from four donors at day 0.**

The UMAP visualizations of cells colored by gene signature scores computed by Gene set variation analysis (GSVA) for donor 1 **(a)**, donor 2 **(b)**, donor 3 **(c),** and donor 4 **(d)** on M-CSF stimulation CD14 monocytes scRNA-seq dataset, day0.

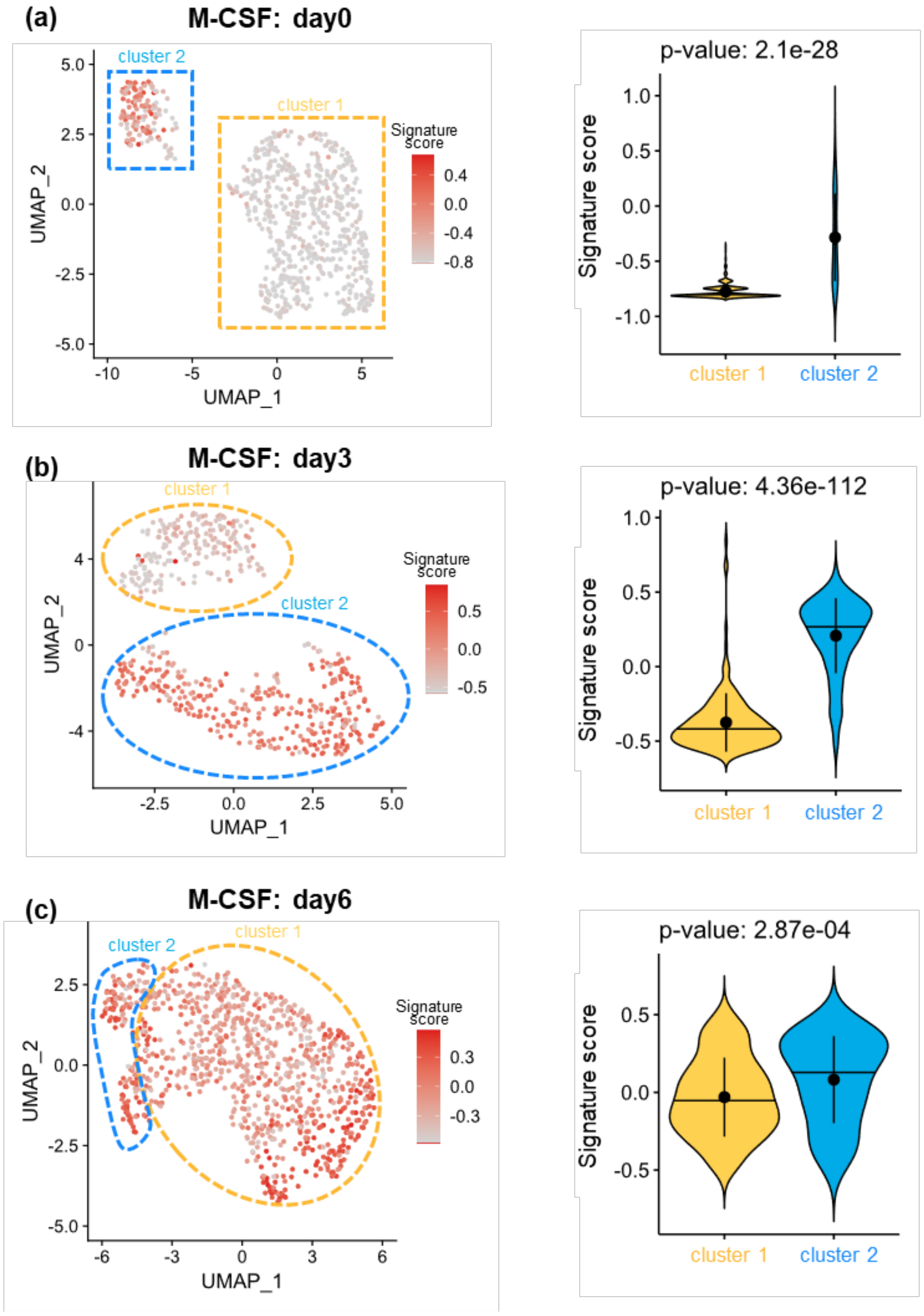

**Supplementary Fig. 16 Results on M-CSF stimulating CD14 monocytes scRNA-seq dataset at day 0, 3 and 6 for donor 2.**

The UMAP visualizations of cells colored by gene signature scores for donor 2 on M-CSF stimulation CD14 monocytes scRNA-seq dataset at day0 **(a)**, day3 **(b),** and day6 **(c)**.

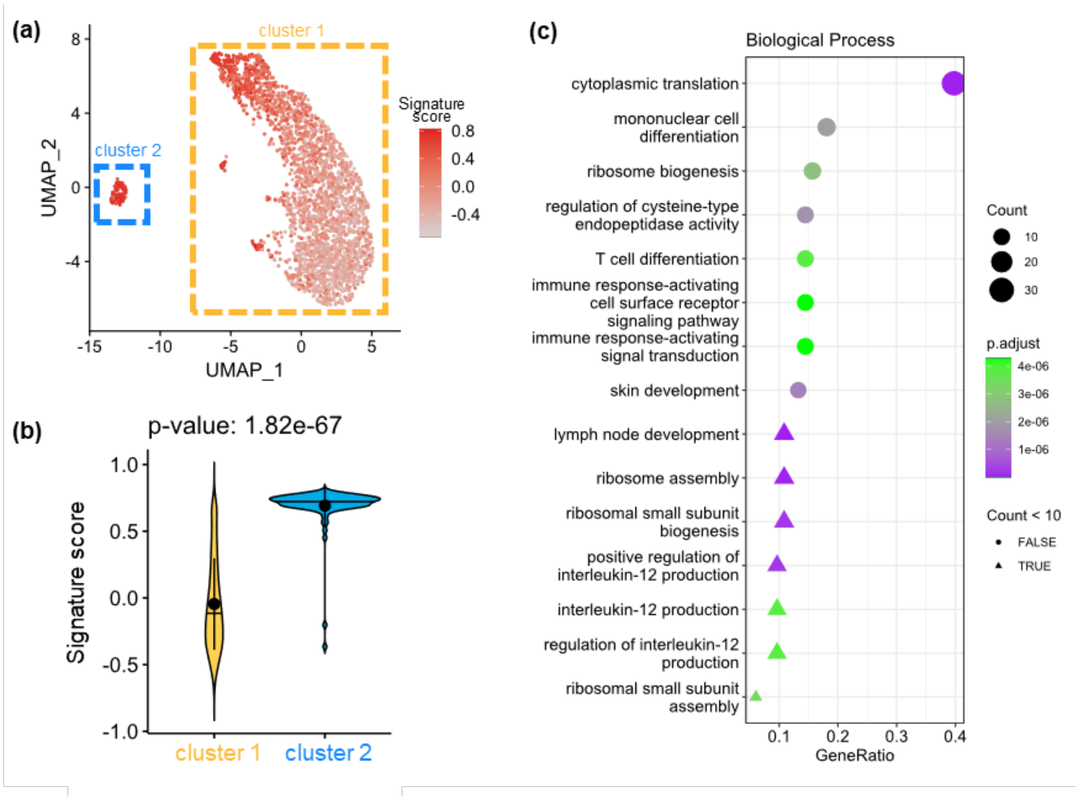

**Supplementary Fig. 17 Results on the 10X Genomics CD14 monocytes scRNA-seq dataset.** **(a)** The UMAP visualizations of cells colored by gene signature scores for 10X Genomics CD14 monocytes scRNA-seq dataset. **(b)** The violin plots show the difference in signature scores between the two cell clusters. **(c)** The top fifteen biological processes involved in the gene signature, implemented by the R package “clusterProfiler’’, using all genes in this dataset as the background gene set.

### 6. Supporting results for main Fig.4

In this section, we will give some details on how SCARP discovered key factors of melanoma by portraying co-accessibility relationships on the SOX10KD scATAC-seq dataset.

We first annotated the types of accessible regions, such as introns and promoters, in the SOX10KD scATAC-seq data. As shown in the upset plot in Supplementary Fig. 18a, a region may be annotated to more than one type, we used the default priority in the R package 'ChIPseeker' to make unique assignments (promoter >= 5'UTR >= 3'UTR >= Exon >= Intron >= Downstream >= Distal intergenic). To further map peaks to genes, we screened those peaks that are annotated as promoters of specific genes (within $\pm$1kb of TSS), and treated them as genes directly, as shown in Supplementary Fig. 19. Besides, to investigate the biological significance of each feature of the peak's low-dimensional representations obtained by SCARP, we measured the enrichment of the different region types within the features by computing the AUC scores, as shown in Supplementary Fig. 20.

The regulatory relationships of the top 50 genes with the highest correlation to SOX10 were validated by the Reactome database. We further performed functional enrichment analysis of the genes in the GRN, and the cnetplot of KEGG analysis results was shown in Supplementary Fig. 21. The gene set GO enrichment analysis, including molecular function, cellular components, and biological process analysis were shown in Supplementary Fig. 22.

Results obtained by scAND, cisTopic, and original methods using the same analysis steps on the SOX10KD dataset as SCARP did, including validation of regulatory relationships (GRN) and gene functional enrichment analysis, are shown in Supplementary Fig. 23, Supplementary Fig. 24, and Supplementary Fig. 25, respectively.

Furthermore, we also performed survival analysis using the R package 'survival' and found that the abnormal expression of some genes was associated with poor prognoses, as shown in Supplementary Fig. 26.

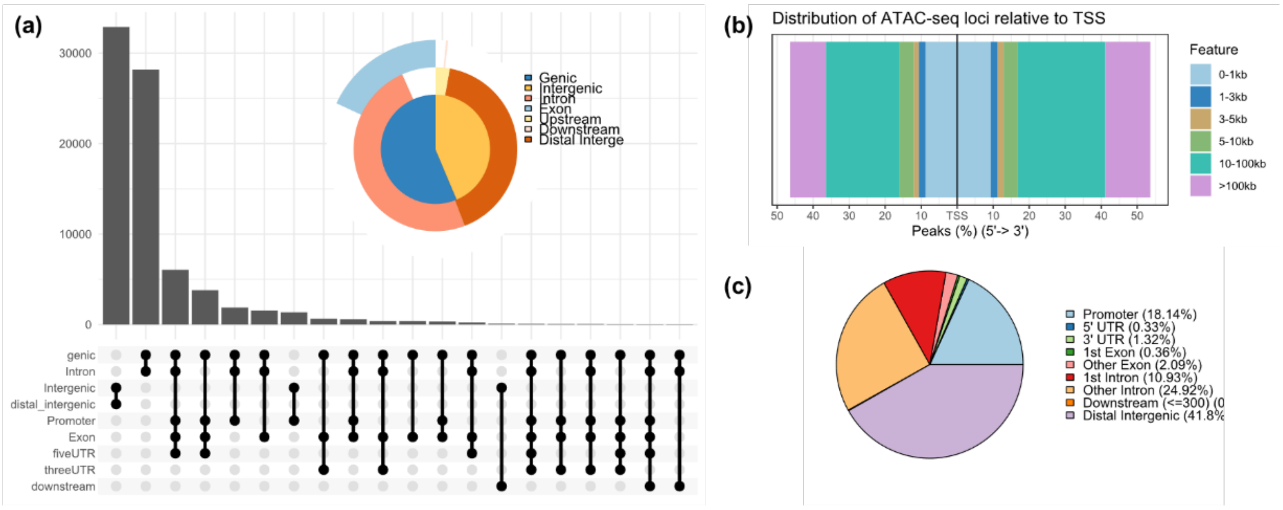

**Supplementary Fig. 18 Peaks annotation result on SOX10KD scATAC-seq dataset.**

**(a)** The upset plot of peaks annotation. To assign a peak to a specific region type, the priority was set as: promoter >= 5’UTR >= 3’UTR >= Exon >= Intron >= Downstream >= Distal intergenic. **(b)** The distance distribution of the loci relative to TSS. **(c)** The proportions of different genome types annotated from peaks.

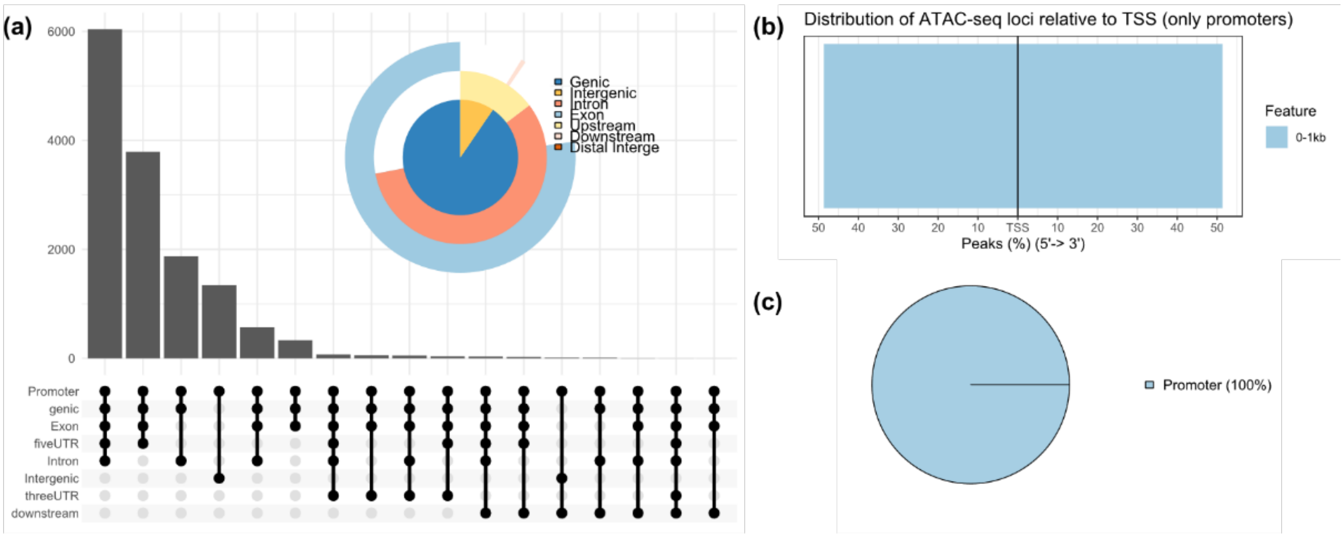

**Supplementary Fig. 19 Same as Supplementary Fig. 18, except that we only selected those peaks annotated as prompters.**

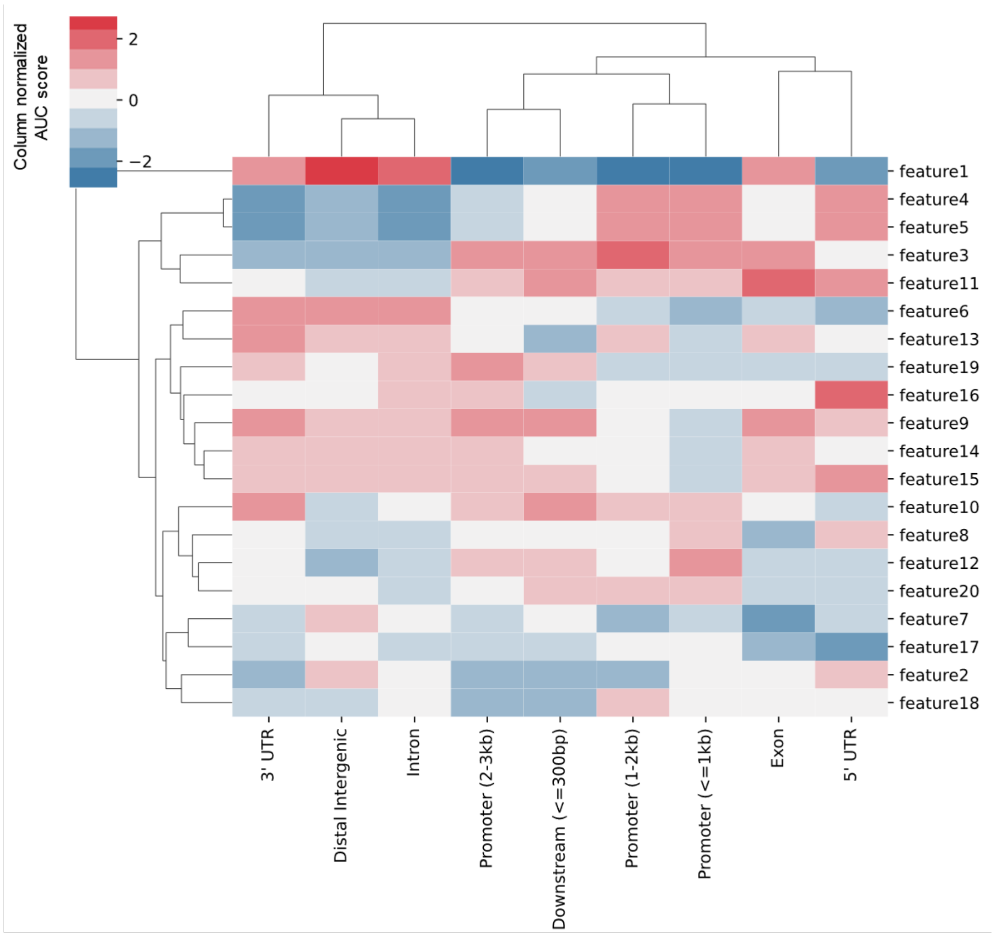

**Supplementary Fig. 20 We measured the enrichment of the different region types (columns) within the features (rows) by computing the AUC scores.** This result demonstrated that the biological significance of SCARP-derived peak low-dimensional features may be related to their region types, since different features showed different region type preferences, and in turn different region types were enriched in different features. We clustered the columns (region types) of this heatmap and found that certain regions with similar regulatory functions had feature preference consistency. For example, proximal promoters (<=1kb & 1-2kb) were similar to each other and showed consistency with 5' UTR, and these regulatory elements were located upstream of the gene and usually initiated gene transcription by binding transcription factor; similar consistency results were found for 3' UTR and intron. This indicated that there were clear differences in accessibility between the different region types, and they were closely related to their regulatory functions.

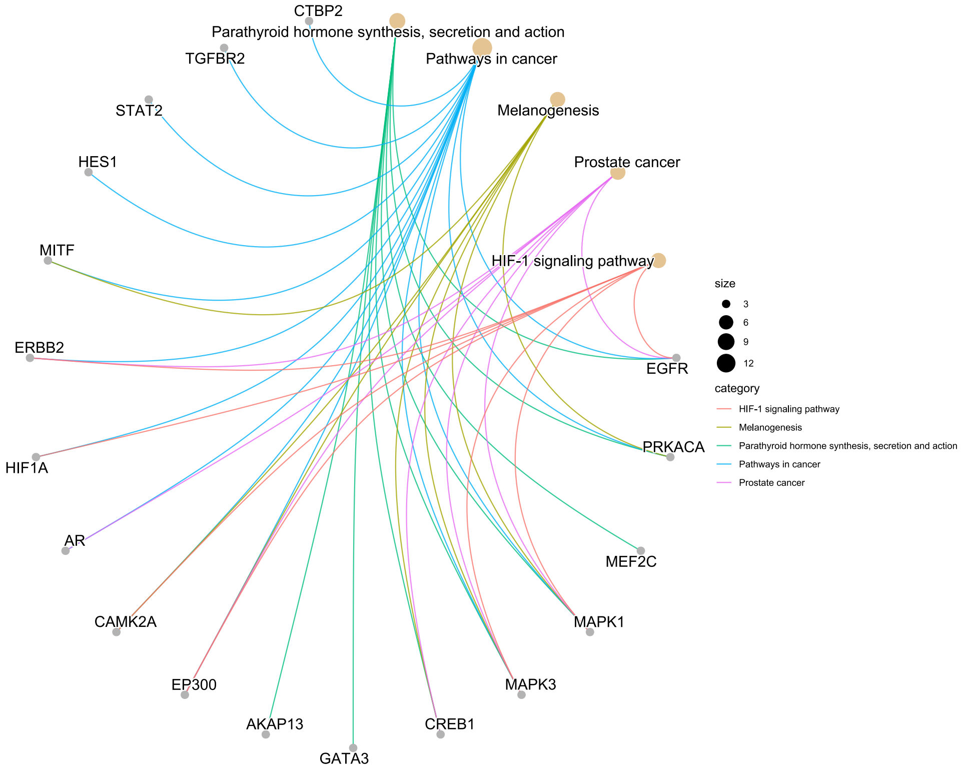

**Supplementary Fig. 21 The cnetplot of KEGG analysis of gene set in the gene regulatory network constructed by inputting genes obtained by SCARP into Cytoscape.**

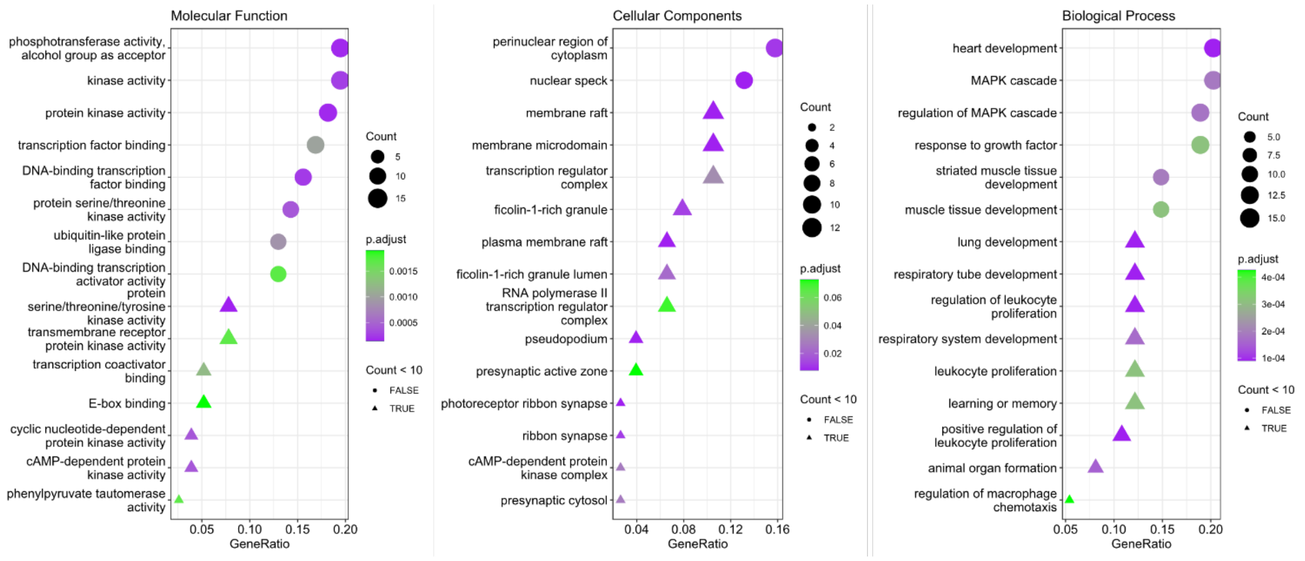

**Supplementary Fig. 22 Gene GO enrichment analysis on the SCARP-derived genes, including molecular function, cellular components, and biological process analysis.**

Details on each GO term was given in Supplementary Table 5.

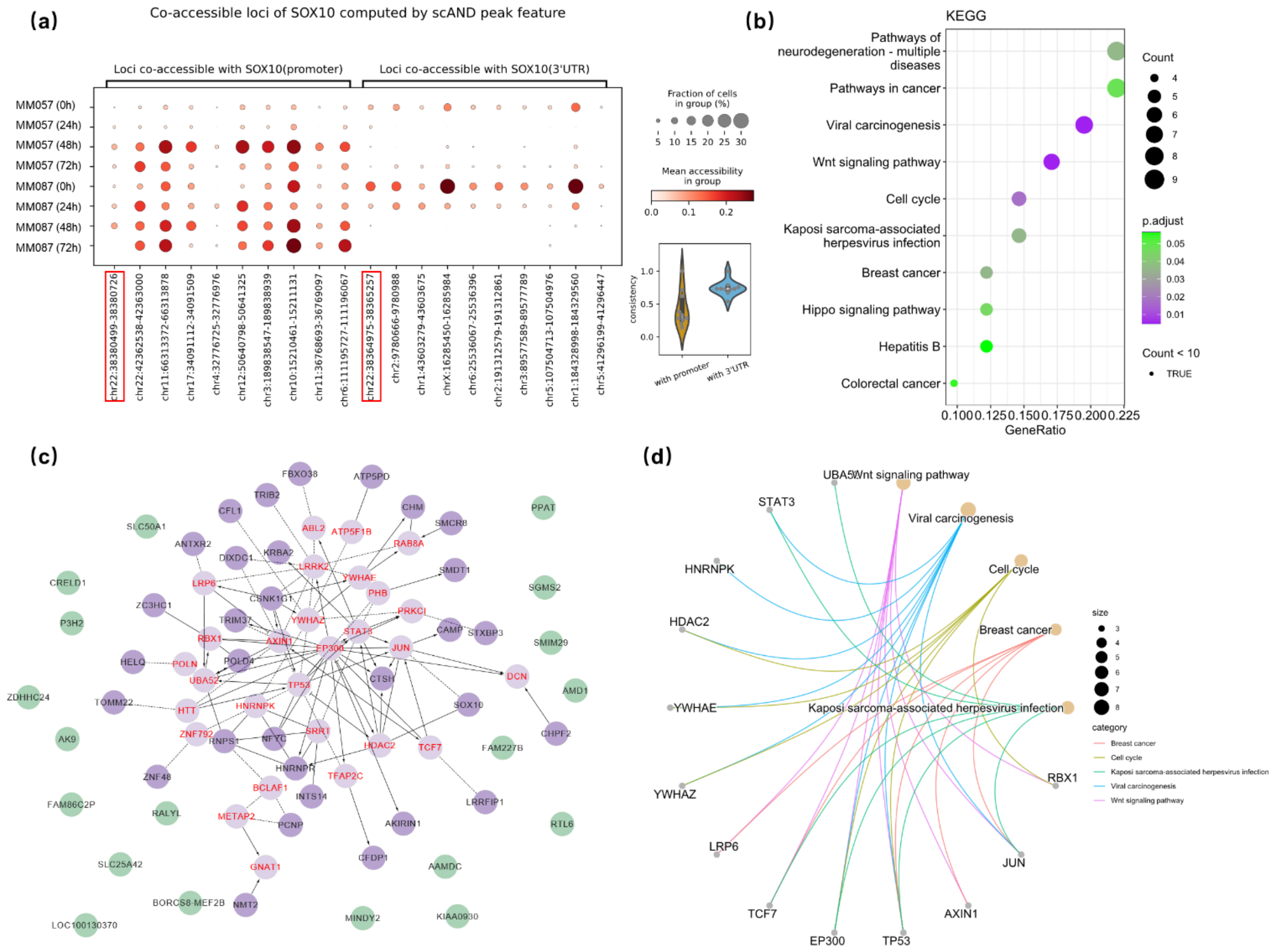

**Supplementary Fig. 23  Results on the SOX10KD dataset using the scAND method.**

**(a)** Changes in accessibility over time for those loci highly associated with the SOX10 promoter and those highly associated with the SOX10 3'UTR. **(b)** The KEGG pathway analysis. **(c)** The GRN outputted by Cytoscape. **(d)** The cnetplot of KEGG analysis.

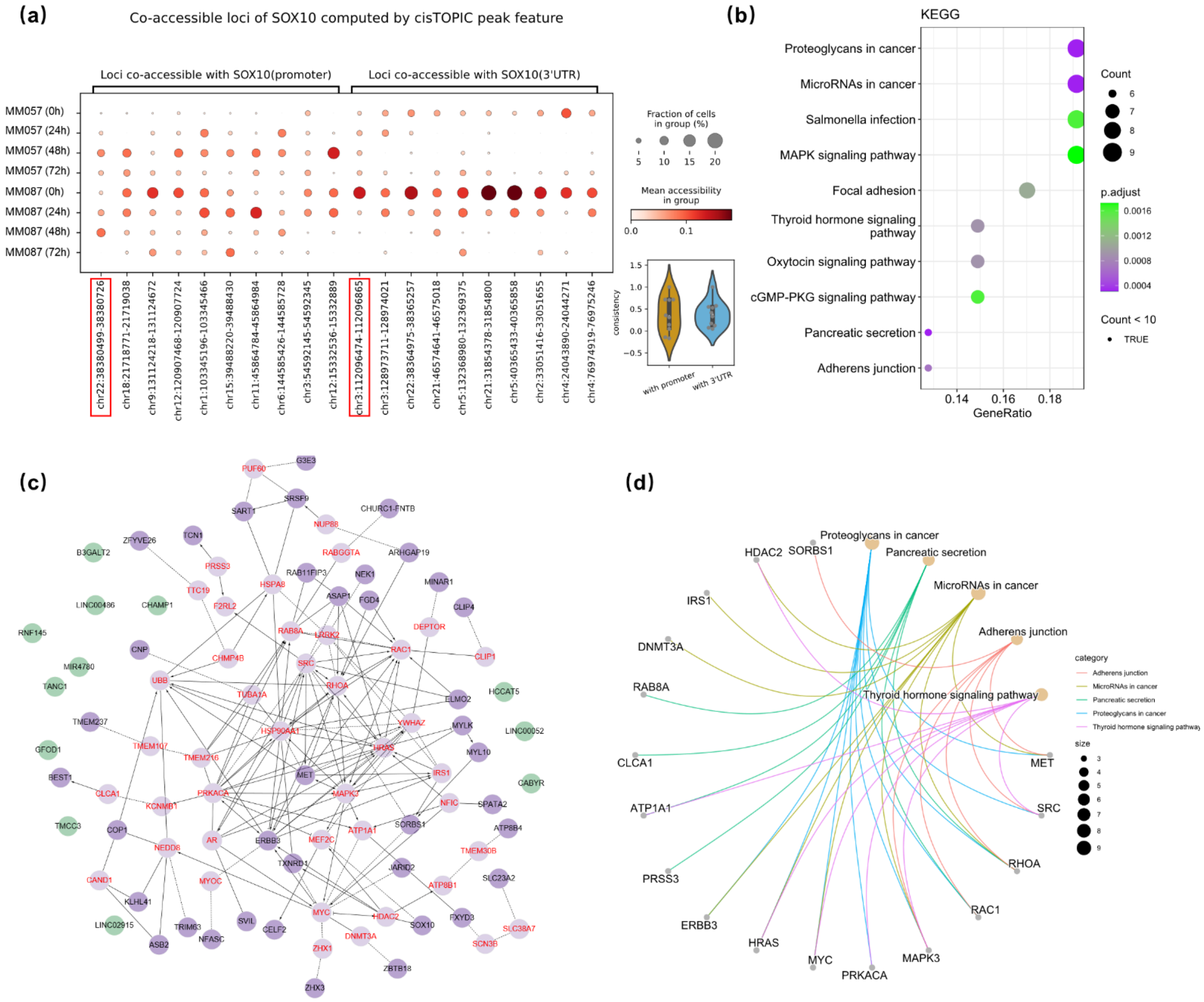

**Supplementary Fig. 24 Results on the SOX10KD dataset using cisTopic method.**

**(a)** Changes in accessibility over time for those loci highly associated with the SOX10 promoter and those highly associated with the SOX10 3'UTR. **(b)** The KEGG pathway analysis. **(c)** The GRN outputted by Cytoscape. **(d)** The cnetplot of KEGG analysis.

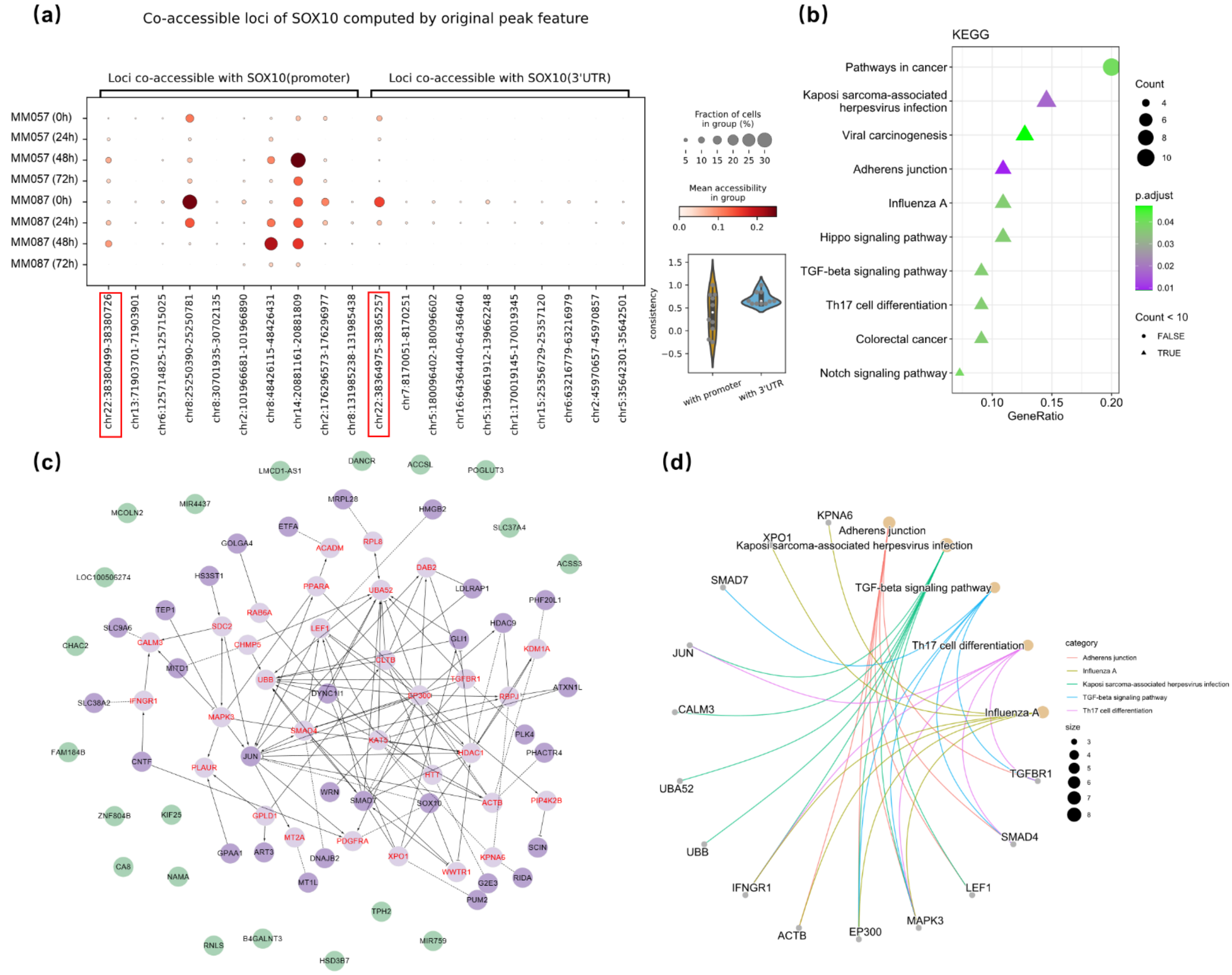

**Supplementary Fig. 25 Results on the SOX10KD dataset using original method.**

**(a)** Changes in accessibility over time for those loci highly associated with the SOX10 promoter and those highly associated with the SOX10 3'UTR. **(b)** The KEGG pathway analysis. **(c)** The GRN outputted by Cytoscape. **(d)** The cnetplot of KEGG analysis.

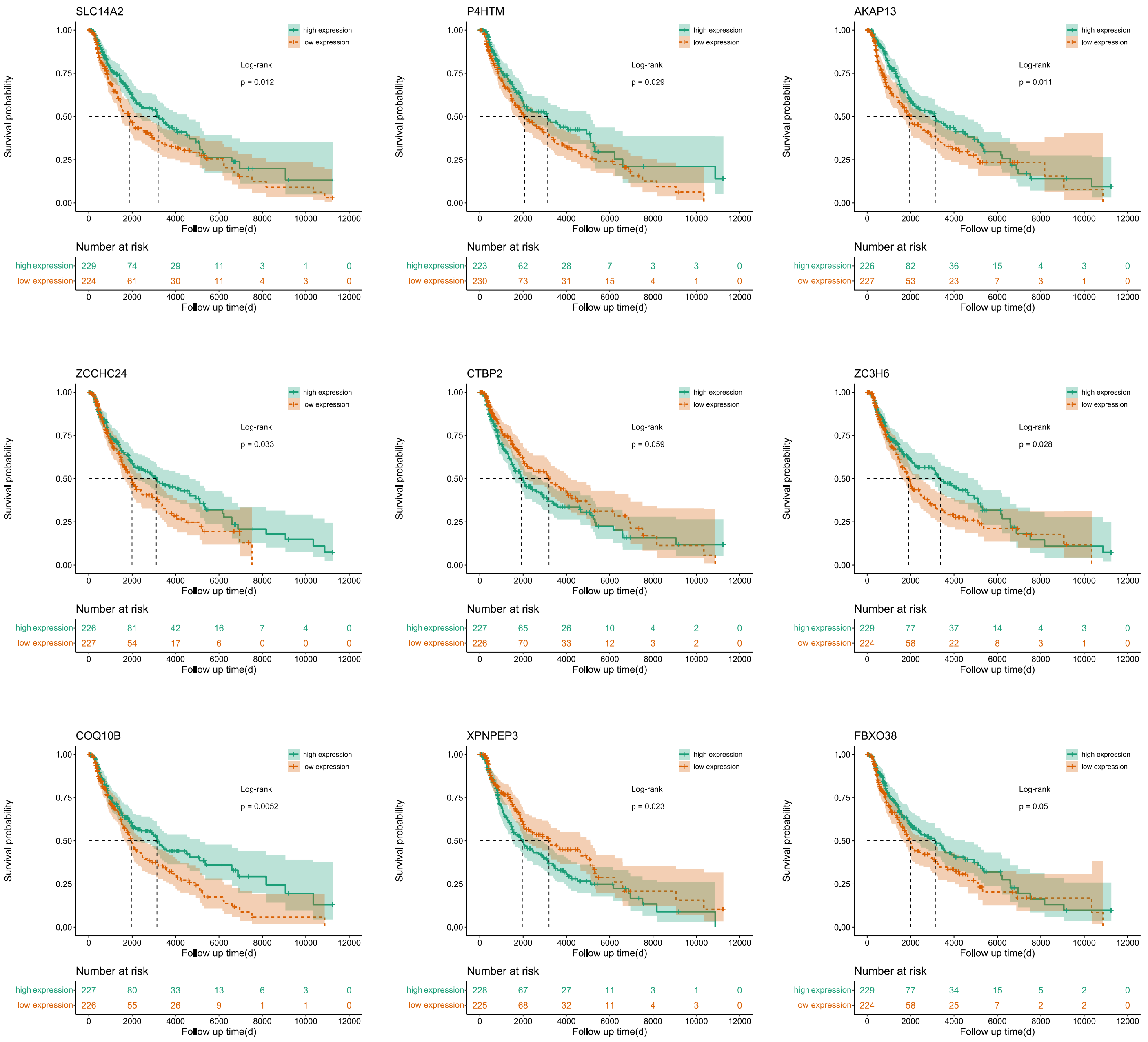

**Supplementary Fig. 26 Survival analysis on some of the SCARP-derived genes.**

Survival analysis results of genes *SLC14A2*, *P4HTM*, *AKAP13*, *ZCCHC24*, *CTBP2*, *ZC3H6*, *COQ10B*, *XPNPEP3*, and *FBXO38*.

### 7. Supporting results for main Fig.5

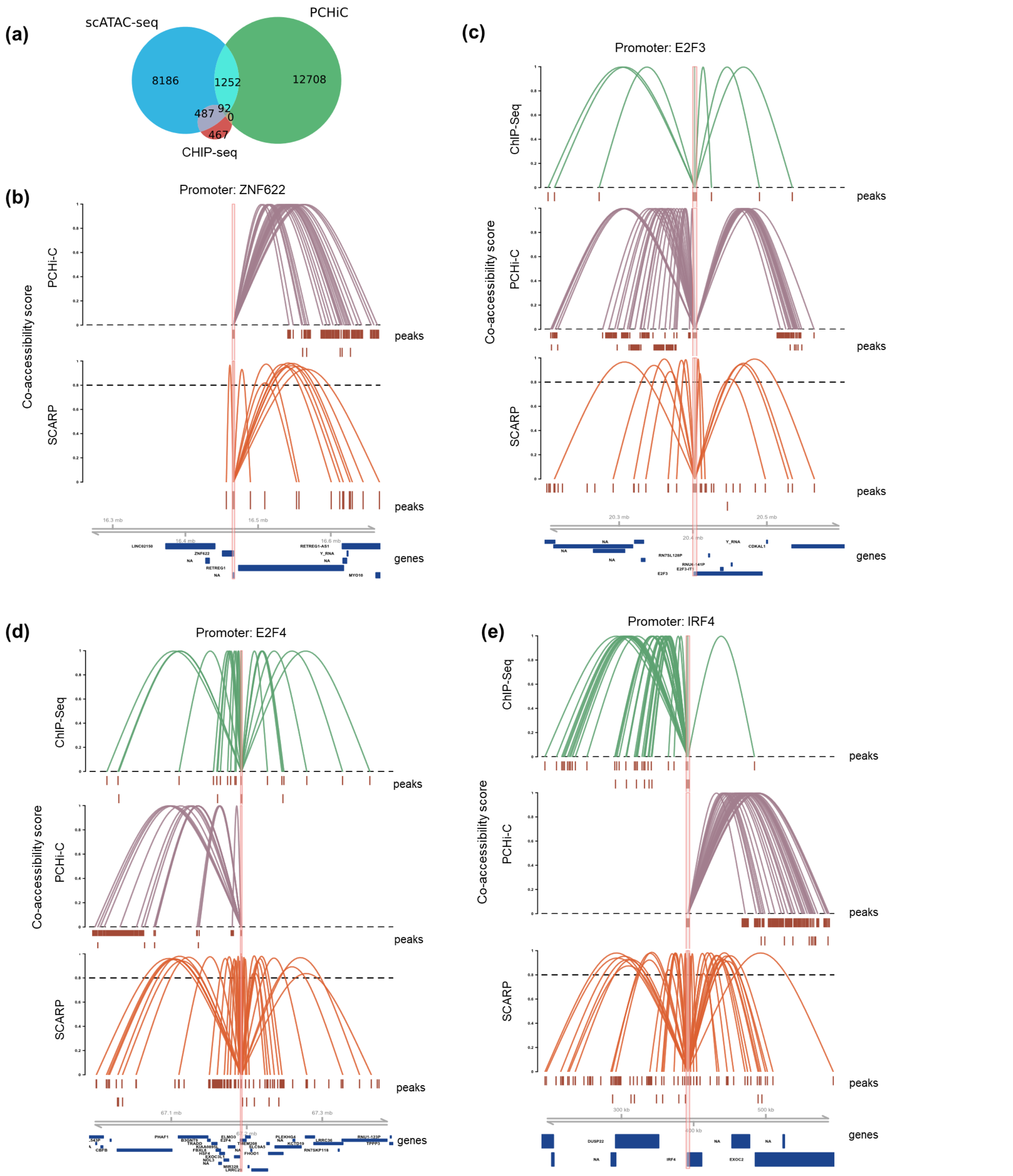

**Supplementary Fig. 27 SCARP-derived cis-regulatory interactions.**

**(a)** The Venn plot of genes in three types of data. (b)(c)(d)(e), SCARP-derived cis-regulatory interactions of genes **(b)** *ZNF622*, **(c)** *E2F3*, **(d)** *E3F4*, and **(e)** *IRF4*, along with external PCHi-C and ChIP-seq evidence. The legends are the same as in Fig. 5e of the main text.

### 8. Rationality of integrating peak location information.

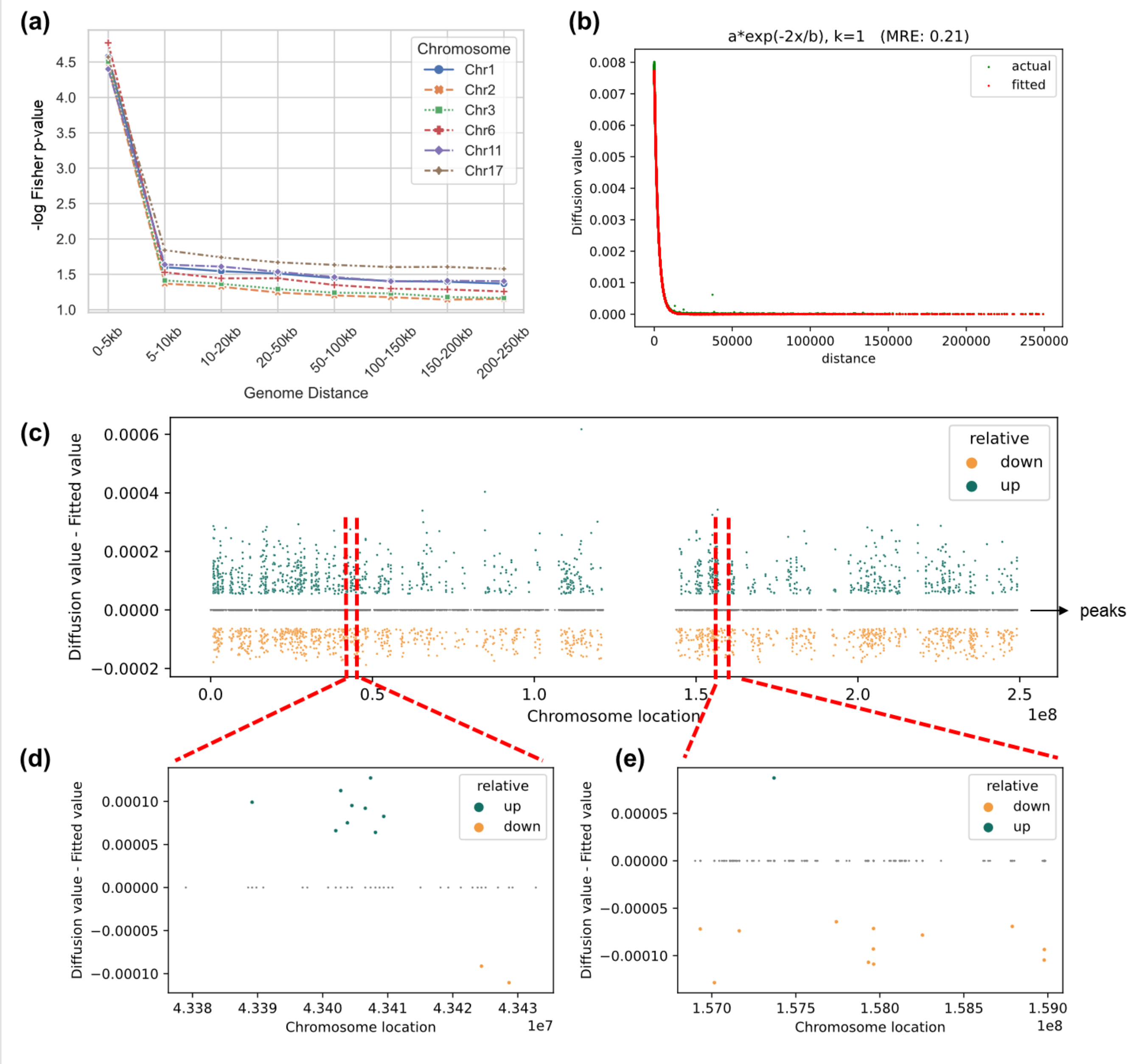

**Supplementary Fig. 28 Elaboration of the rationality of peaks location information.**

**(a)** Peaks closer in the genome are more likely to be co-accessible. **(b)** Taking chromosome 1 of blood2K datasets as an example, we extract the sub-diagonal elements of the NR diffused matrix (green) and fit them (red) with function $a * exp(-2x/b)$. **(c)** The distribution of difference (NR diffusion value minus fitted value) over the genome positions of chromosome 1. We found that those peaks with network weights higher (or lower) than expected after diffusion show a certain pattern on the genome. **(d)** and **(e)** zoom in on a specific region of (c).

### 9. Kept components of SCARP on various datasets in this study.

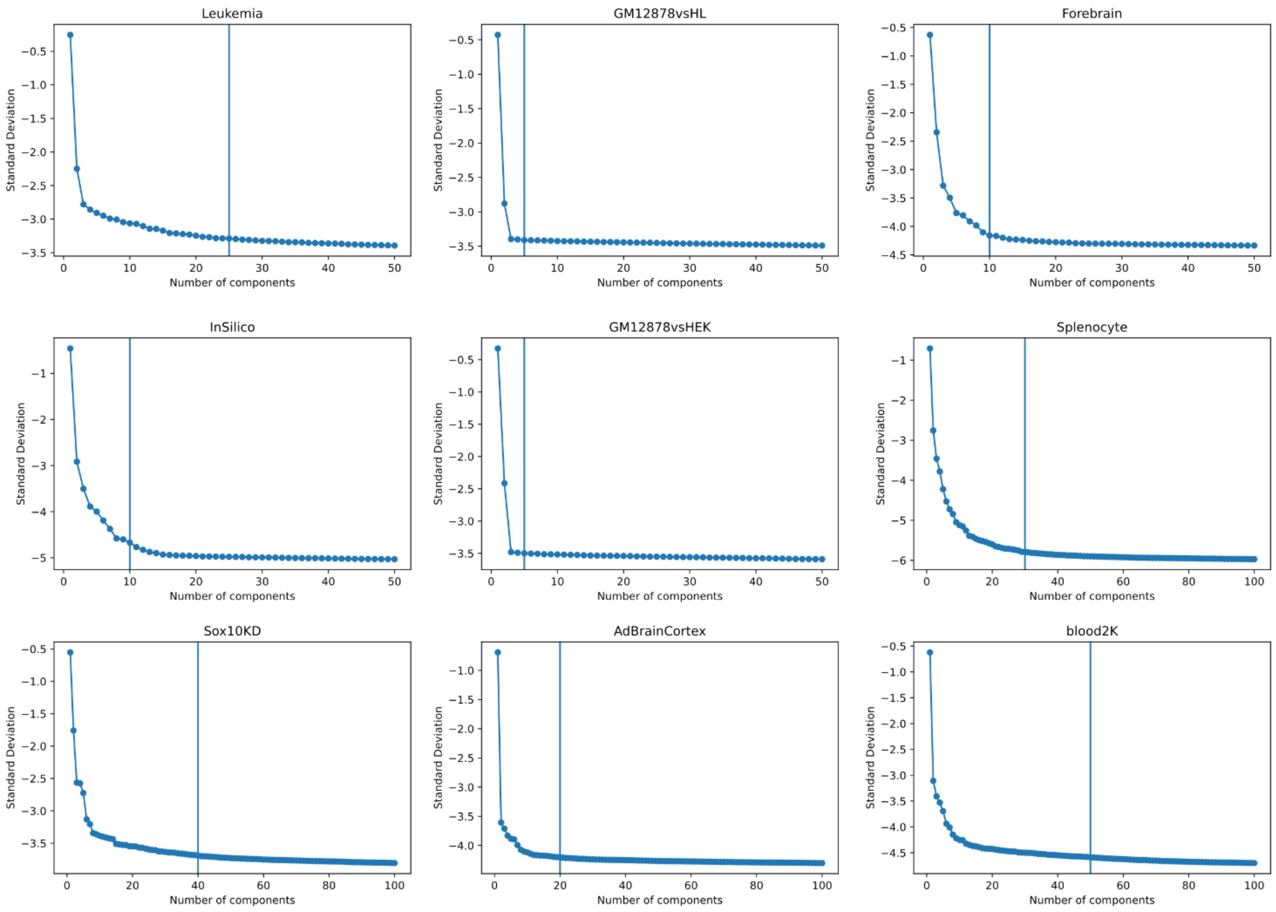

**Supplementary Fig. 29 Number of kept components of SCARP on various scATAC-seq datasets.**
